# When non-canonical olfaction is optimal

**DOI:** 10.1101/2025.02.27.640624

**Authors:** Caitlin Lienkaemper, Meg A. Younger, Gabriel Koch Ocker

## Abstract

The early olfactory system is canonically described by a “one-receptor-to-one-neuron” model: each olfactory sensory neuron (OSN) expresses a single type of olfactory receptor. Although the olfactory systems of many model organisms approximately follow this canonical organization, a number of exceptions are known. In particular, Aedes aegypti mosquitoes co-express multiple types of olfactory receptors in many OSNs. Why do some olfactory systems follow the canonical organization while others violate it? We approach this question from the normative perspective of efficient coding. We find that the canonical and non-canonical organizations optimally encode odor signals in different types of olfactory environment. Non-canonical olfaction is beneficial when relevant sources emit correlated odorants and the environment contains odorants from ethologically irrelevant odor sources. Our theory explains previous observations of receptor co-expression and provides a framework from which to understand the structure of early olfactory systems.

---

In model organisms such as mice and fruit flies, the early olfactory system primarily follows a canonical “one-receptor-to-one-neuron” organization (Fig. 1A, top). That is, each olfactory sensory neuron (OSN) expresses a single type of olfactory receptor (OR) [1–4] and neurons that express the same OR project to a dedicated glomerulus or glomeruli, located in the olfactory bulb in mammals [5–8] and the antennal lobe in insects [3, 9, 10]. This allows downstream regions in the brain to decode odor identity and concentration from the interaction of each odorant with each type of olfactory receptor [1, 11].

**Fig. 1.**
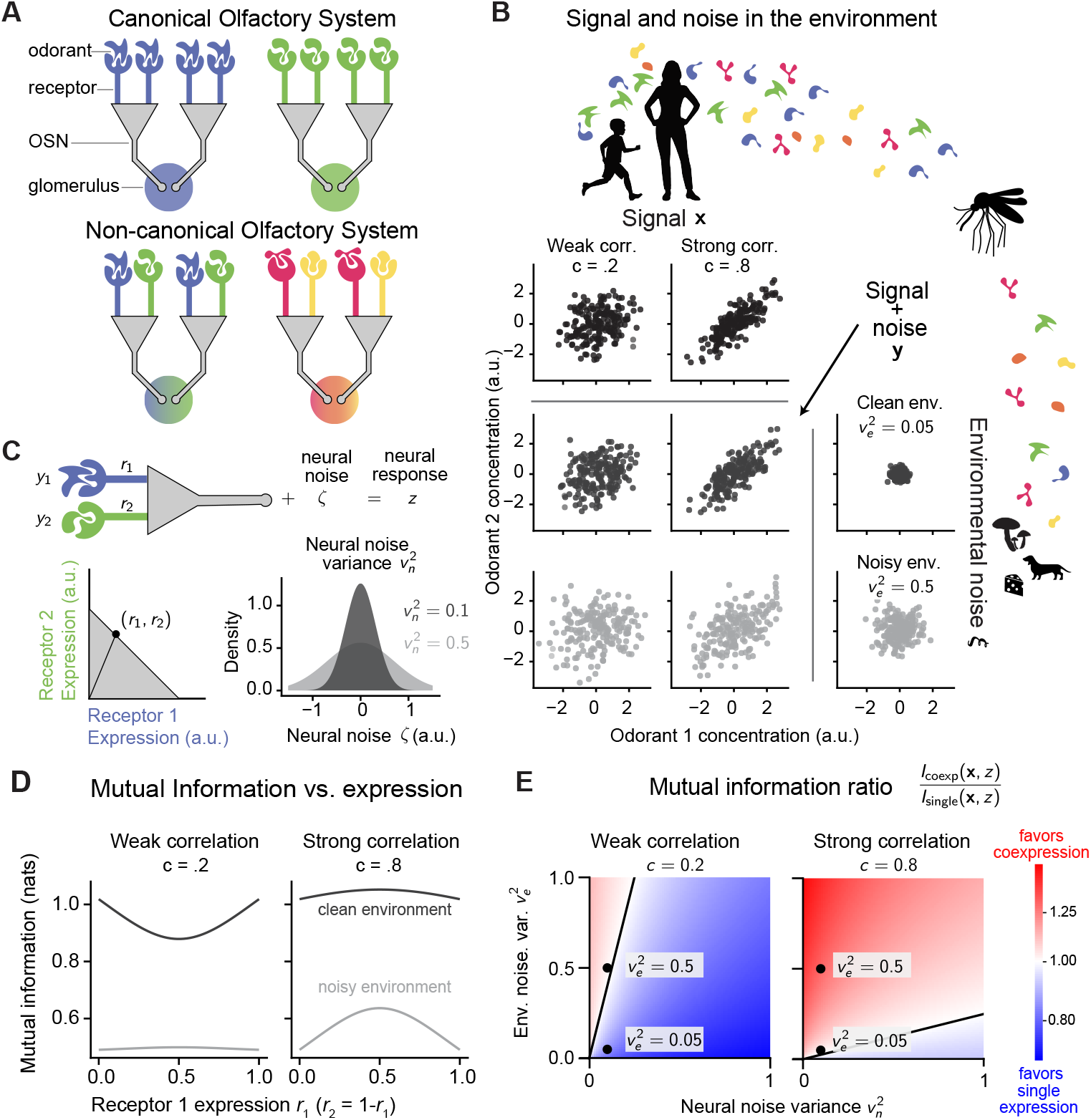
Co-expressing receptors for odorants which are correlated in the environment maximizes mutual information between olfactory signal and neural response in conditions of high environmental noise. (A) Canonical (top) and non-canonical (bottom) early olfactory systems. (B) Olfactory stimuli, i.e., odorant concentrations at the mosquito antenna, consist of both signal and noise. (Top) The signal components derive from ethologically relevant sources. (Right) Odorants are regarded as noise if they do not arise from an ethologically relevant source. (C) We model the neural responses as a noisy sum of odorant concentrations, weighted by receptor expression levels. We model response noise as independent across neurons, and denote its variance with the parameter *v*_*n*_^2^. (D) Mutual information between model OSN response and odorant signal, as a function of receptor expression levels. Solid line: clean environment (*v*_*e*_^2^ = .05). Shaded line: noisy environment (*v*_*e*_^2^ = .5). Response noise level: *v*_*n*_^2^ = .1. (E) Mutual information ratio 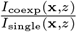 as a function of *v*_*e*_^2^ and *v*_*n*_^2^. *I*_coexp_(**x**, *z*) is the mutual information between olfactory signal and the response of a neuron which co-expresses both receptors equally, and *I*_single_(**x**, *z*) is the mutual information between olfactory signal and the response of a neuron which single-expresses receptor one. Black line marks the boundary between the region where single expression is optimal and the region where co-expression is optimal.

However, a number of violations of the canonical model have been observed across diverse species (Fig. 1A, bottom) [12–19]. Here, we focus on one: the *Aedes aegypti* mosquito. *Ae. aegypti* express many types of chemoreceptors (102 - 178, depending on expression threshold) relative to the number of glomeruli they have (roughly 65) [20–22]. Single nucleus RNA sequencing indicates widespread co-expression, with OSNs co-expressing up to six OR types [22, 23]. The discovery of widespread co-expression in *Ae. aegypti*, and beyond, invites us to ask why some olfactory systems obey the canonical model while others do not.

*Ae. aegypti* are highly effective olfactory predators, relying on chemosensory cues including CO_2_ from breath and volatile components of human body odor to locate human hosts for blood meals. In addition, *Ae. aegypti* use olfactory cues to find floral resources for nectar feeding [24], mates [25], and oviposition sites [26]. Experiments inactivating receptor families show that the *Ae. aegypti* olfactory system is extremely robust. Mutant mosquitoes with many non-functional ORs maintain attraction to humans and the ability to locate humans in naturalistic settings [27–30].

The *Ae. aegypti* olfactory system must also, like all olfactory systems, be robust to naturalistic sources of noise in neural responses and in the environment [31, 32]. The ability of chemosensory neurons, such as OSNs, to detect the local concentration of an odorant is limited by thermodynamic noise, i.e., the fact that odorants arrive at the receptor at random rates [33, 34]. For example, electrical noise caused by the stochastic opening and closing of ion channels leads to fluctuations in membrane potentials; this can lead to additional variability in stimulus responses [35–38]. *Ae. aegypti* locate humans using olfactory cues that have diffused over long distances and in the presence of interference from odorants emitted by irrelevant odor sources. Together, this suggests that the robustness of *Ae. aegypti* olfactory behavior may be related to their non-canonical early olfactory system.

In this paper, we aim to answer the following questions: what fitness advantage, if any, does OR co-expression confer to *Ae. aegypti* and other species that exhibit it? What fitness advantage does the canonical organization confer to species which follow it? If co-expression is advantageous, which receptors should be co-expressed?

We approach these questions from the perspective of the efficient coding hypothesis, which states that sensory systems adapt to the structure and statistics of natural stimuli to maximize information transmission while minimizing resource costs [39–41]. Specifically, we hypothesize that OR co-expression patterns maximize the information OSN activity carries about behaviorally relevant odorant mixtures, under a constraint on the total amount of receptor in each neuron. This implies that OR co-expression and canonical expression confer the same fitness advantage (maximizing OSN response information) under different circumstances.

Through simulations and analytical calculations, we show that three factors determine whether the optimal expression patterns will be canonical or co-expressing: the level of noise in the olfactory environment, the level of noise in neural responses, and the correlation of odorants across ethologically relevant stimuli. We propose that receptors should be co-expressed when they bind odorants that are reliably co-emitted by behaviorally relevant sources, and environmental fluctuations in odorant concentrations are the dominant source of noise in neural activity. These predictions are confirmed by the affinities of co-expressed receptors in *D. melanogaster*. Finally, we show that when co-expression or single-expression are optimal, they endow an OSN with different sensory computations: detecting principal components of the input or maximizing response gain.

## 1 Results

### 1.1 Optimal receptor expression patterns in a single OSN

Our goal is to understand how the structure of behaviorally relevant stimuli and the sources of noise affect whether or not receptor co-expression is optimal. We first address this question using the simplest model which incorporates all of these features, a single OSN in an environment with one relevant olfactory source (Fig. 1B,C). Our first model of an OSN has two ORs in its genome, expressed at the relative levels *r*_1_ and *r*_2_. A canonical OSN would have either (*r*_1_, *r*_2_) = (1, 0) or (0, 1), while a non-canonical OSN would have intermediate expression levels (Fig. 1A,C).

The olfactory environment contains an ethologically relevant odor source emitting two odorants with concentrations **x** = (*x*_1_, *x*_2_). Before the odorants reach the OSN, their relative concentrations can change due to diffusion or other, behaviorally irrelevant, sources of the same odorants (Fig. 1B, right). Together, the odor that reaches the OSN has the relative concentrations

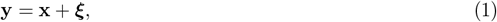

where **y** = (*y*_1_, *y*_2_) are the odorant concentrations at the OSN, **x** = (*x*_1_, *x*_2_) are the odorant concentrations at the source and ***ξ*** = (*ξ*_1_, *ξ*_2_) is the environmental noise in the odorant concentrations. Since the two odorants are emitted by the relevant common source, their concentrations are correlated with correlation coefficient *c* (Fig. 1B, top). We parameterize the noise level of the odorants with the variance *v*_*e*_^2^, which could correspond to the OSN’s distance from the ethologically relevant source and/or the prevalence of distractor sources.

According to the efficient coding hypothesis, the OSN should encode the odorant concentrations in the signal as well as possible under its resource limitations [39, 40]. We consider a constraint on the total amount of OR in the neuron: the OSN has a bounded total amount of receptor and cannot obtain more total receptor by co-expression, *r*_1_ + *r*_2_ ≤ 1 (Fig. 1C).

To calculate the information that the neural response carries about the odor concentration at the source, we begin with several simplifying assumptions that we will later relax: (1) the variance of the two odorants at the source are equal; (2) each receptor binds to only one odorant; (3) the neural response *z* is a noisy sum of the odorant concentrations, weighted by the receptor expression levels:

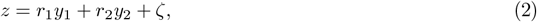

where *ζ* is additive neural response noise (Fig. 1C). The response variability is parameterized by the variance of *ζ, v*_*n*_^2^. We also assume that (4) the signal odor concentrations, noise odor concentrations, and neural response noise follow Gaussian distributions.

The response information depends on 1) the odorant correlation at the relevant source, *c*; 2) the environmental noise level *v*_*e*_^2^; 3) the neural response noise level *v*_*n*_^2^; and 4) the expression levels (*r*_1_, *r*_2_). Depending on these factors, the response information can be maximized by a single-expressing OSN or a co-expressing OSN (Fig. 1D). We computed the optimal expression pattern (*r*_1_, *r*_2_) as a function of the other parameters (Methods 3.1, 3.2). The optimal expression pattern is non-canonical when

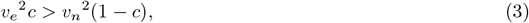

when the quantity *c/*(1 − *c*), which increases as the odorants become more correlated, is greater than the ratio of the response variability and environment noise level, *v*_*n*_^2^*/v*_*e*_^2^. Conversely, the canonical one-receptor-to-one-neuron pattern is optimal when *v*_*e*_^2^*c < v*_*n*_^2^(1 − *c*) (Fig. 1E). This implies that when odorants are uncorrelated across samples of the relevant source (*c* = 0) or the environment is noiseless (*v*_*e*_^2^ = 0), canonical single-expression is always optimal. Likewise, single expression is always optimal when odorants are negatively correlated in the relevant source. When odorants are perfectly correlated (*c* = 1), co-expression is always optimal.

Different odorants’ concentrations may have different levels of variance across natural olfactory scenes (Fig. 2A,B). Therefore, we first relax the assumption that the variance of the two odorants at the source are equal. We label the lower-variance odorant as “odorant one”, and the higher variance odorant as “odorant two”, letting 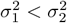 denote the source variances of odorants one and two respectively. Then co-expression is optimal when (Methods 3.3)

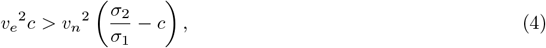

while canonical single expression is optimal when *v*_*e*_^2^*c* ≤*v*_*n*_^2^(*σ*_2_*/σ*_1_ − *c*) (Fig. 2C). Unlike the case where the two source variances are equal, the optimal expression pattern is not symmetric, with higher expression of the receptor for the higher-variance odorant.

**Fig. 2.**
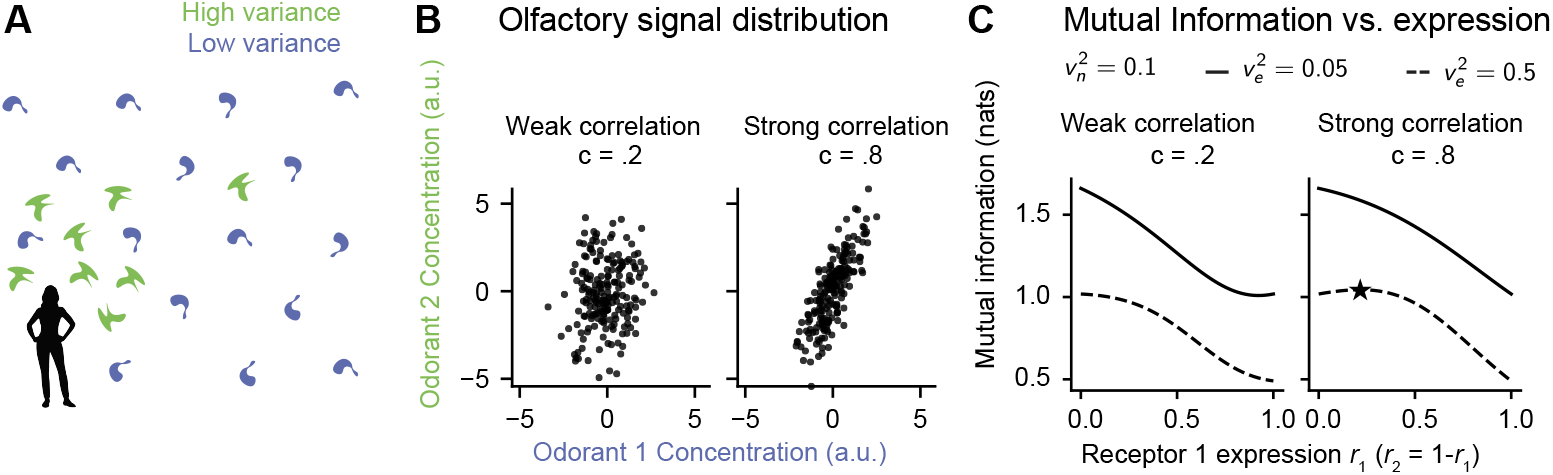
Optimal co-expression favors more expression of the receptor that detects the higher-variance odorant. (A) High vs low variance in odorant concentrations can be due to differences in spatial concentration. (B) Stimulus distributions for weakly correlated stimuli (*c* = .2, left) and strongly correlated stimuli (*c* = .8, right). In both plots, odorant 2 has higher variance, with 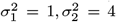. (C) Mutual information as a function of receptor expression levels. When co-expression is optimal (*c* = .8, *v*_*e*_ = .5, optimal expression is marked with a star).

### 1.2 Impact of overlapping receptor affinities

Olfactory receptors typically bind to multiple odorants [42]. We thus relax the assumption that each receptor binds to only one odorant, allowing an overlap *a* between the two receptors’ affinities so that receptor 1 binds to odorant 1 with affinity 1 −*a* and odorant 1 with affinity *a*, and vice versa for receptor 2 (Fig. 3A). Thus, *a* corresponds to the overlap in the two receptors’ affinities: when *a* = 1*/*2 they overlap perfectly and if *a* = 0 or *a* = 1, each receptor responds to only one odorant. There are now two mechanisms that correlate the receptor activations: correlation between odorant concentrations in the olfactory environment and correlation induced by overlapping receptor affinity. Receptors with overlapping receptor affinity can have positively correlated activations even when the odorant concentrations are uncorrelated or even negatively correlated (Fig. 3B,C). Does this additional source of correlation change the optimal receptor expression patterns?

**Fig. 3.**
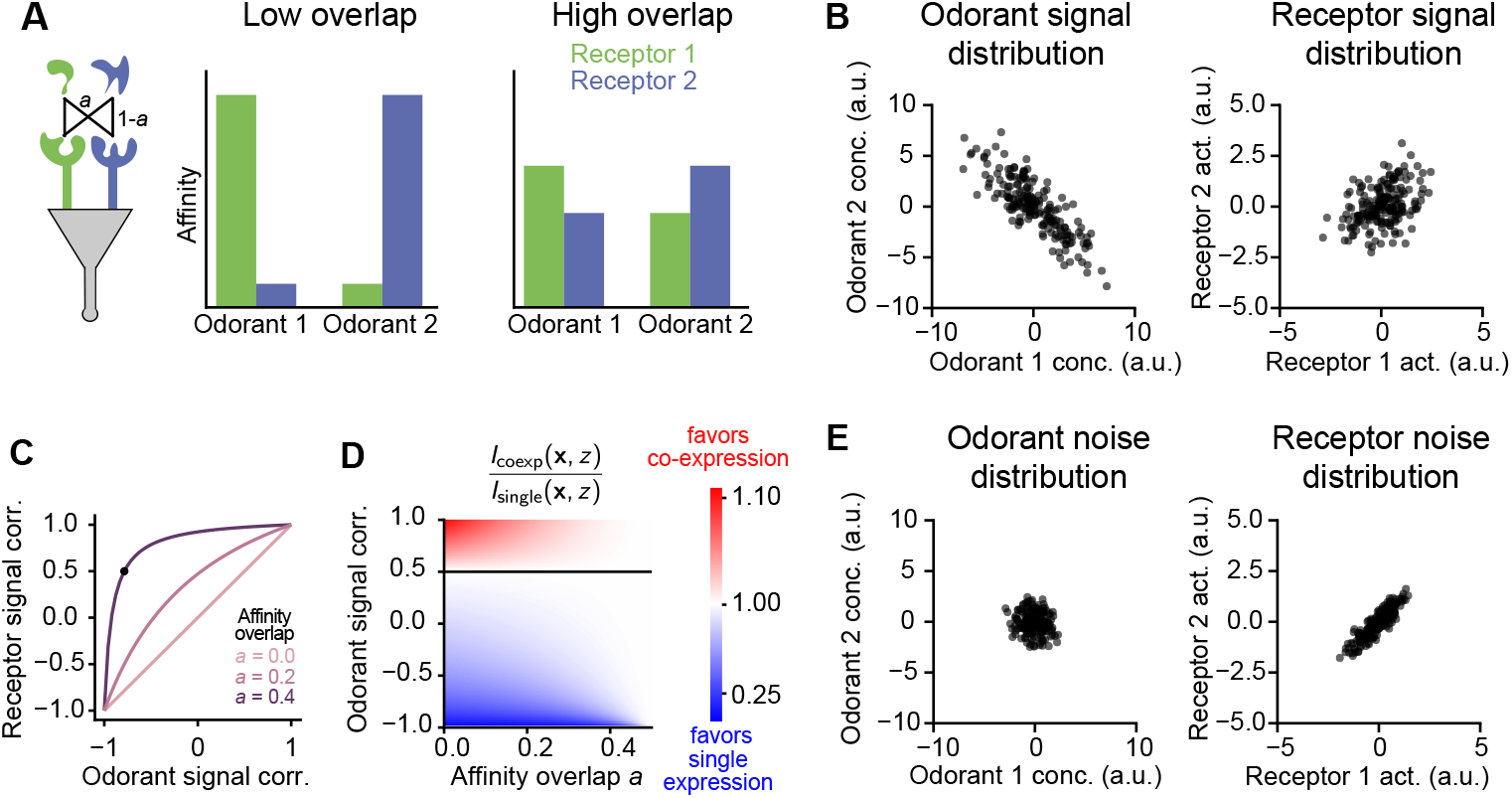
Odorant co-occurrence, rather than receptor affinity overlap, must drive receptor correlations in order for co-expression to maximize mutual information. (A) Low vs high overlap in receptor affinities. (B) Affinity overlap leads to positive receptor signal correlations. Odorant signal correlation *c* = -.79, affinity overlap *a* = 0.4. (C) Receptor signal correlation as a function of odorant signal correlation for various values of the affinity overlap *a*. (D) Mutual information ratio (equal co-expression / single expression) as a function of affinity overlap *a* and odorant signal correlation. Environmental noise variance and neural noise variance are fixed at *ve*2 = .5, *vn*^2^ = .05 respectively. Black line marks the boundary between the region where co-expression is optimal and the region where single expression is optimal. (E) Receptor affinity also leads to positive receptor noise correlations. Odorant signal correlation and receptor affinity overlap as in (B).

We calculated the mutual information between the odorant signal concentrations and the OSN response in the presence of overlapping receptor affinities and determined which receptor expression patterns maximize the information (Methods 3.4). The condition for optimal co-expression is unaffected by the amount of receptor affinity overlap *a*: Eq. 3 still describes when it is optimal to co-express. Rather, overlapping receptor affinities makes the mutual information less sensitive to the expression levels (Fig. 3D). If the two receptors have the same affinity, *a* = 1*/*2, the OSN response does not depend on whether one or the other, or both, are expressed.

Why do overlapping receptor affinities not drive or suppress co-expression? The mutual information can be written in terms of a generalized signal-to-noise ratio involving signal correlations and noise correlations of the neural response (Methods 3.1, Eq. 9). This allows us to write the mutual information in terms of the signal and noise correlations of the receptor activations (Methods 3.4, Eq. 24). We can then express whether or not co-expression is optimal in terms of the receptor signal and noise correlations, *c*^signal^ and *c*^noise^, and common noise variance *σ*^noise^. Co-expression is optimal when

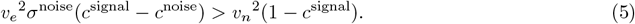

Overlapping receptor affinities drive both response signal correlations and response noise correlations (Fig. 3B). For the affinity matrix 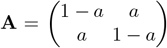, these competing effects cancel. It is thus optimal to co-express receptors when correlated receptor activation is driven by odorant correlations in the olfactory signal, rather than overlapping affinities.

### 1.3 Impact of non-Gaussian odorant distributions

In the previous sections, we assumed that the odorant concentrations in the relevant sources, the environmental noise, and neural response noise all followed Gaussian distributions for the sake of analytical tractability. We now relax these assumptions, and instead study how a more realistic distribution for olfactory signal and noise affect the optimal (co)-expression pattern. We initially address this problem for a single OSN and a pair of receptors which may be co-expressed or not, asking whether the predictions of Equations 3 & 4 remain accurate.

In this section, we model olfactory stimuli as generated from a mixture of odor sources, each of which has a characteristic odor blend that is sparse relative to the total number of odorants (Fig. 4A, Methods 3.5). This results in the distribution of concentrations for each odor having a “spike and slab” structure (Fig. 4B). We designate some odor sources as the olfactory signal and other odor sources as environmental olfactory noise. We again use *v*_*e*_^2^ to denote the variance of the environmental noise, averaged across odorants.

**Fig. 4.**
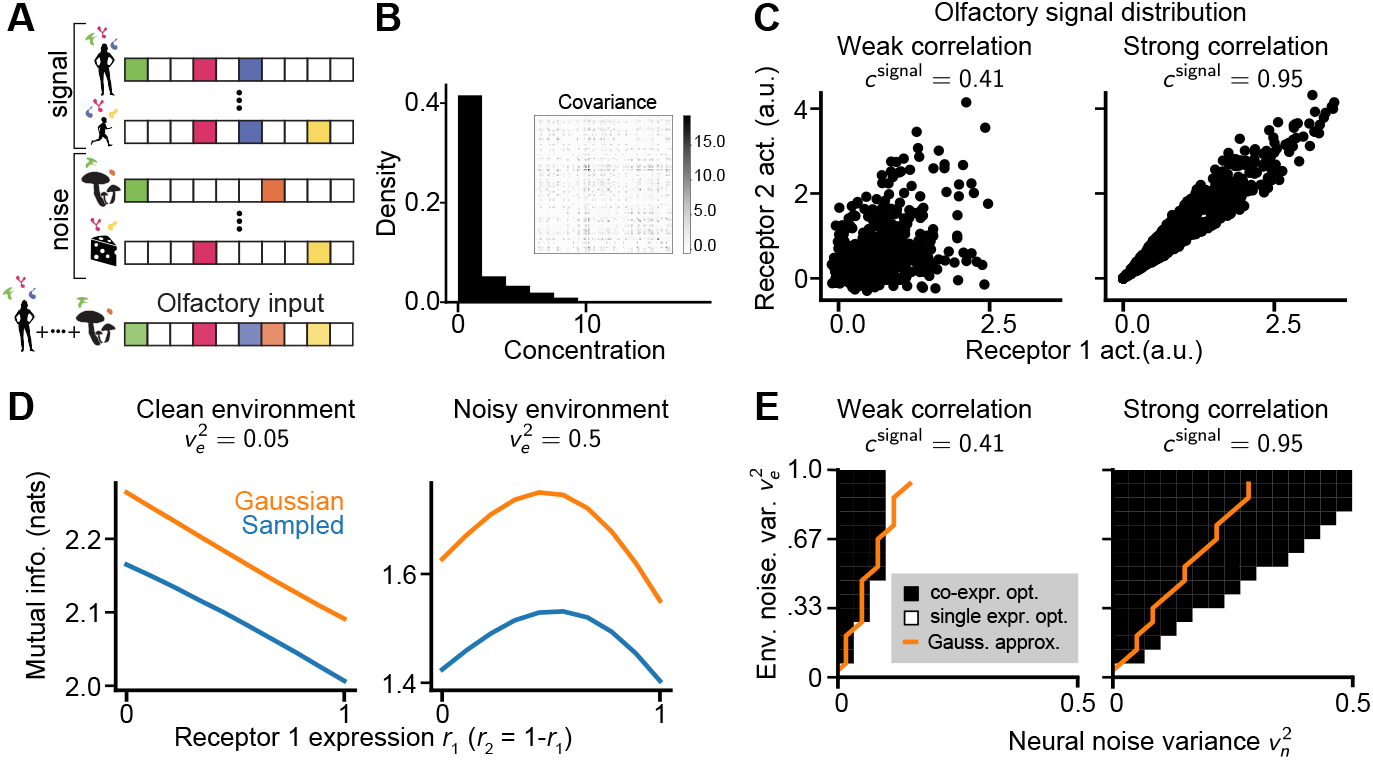
Optimal receptor expression for sparse stimuli is well-approximated by a Gaussian model. (A) Structure of the olfactory stimulus. Top, each odor sources has a characteristic odor blend which is sparse in the space of odors. Bottom, each olfactory stimulus is a sparse mixture of odor sources. (B) Histogram of odor concentrations over all samples in the olfactory signal, for all odorants. Inset: odorant covariance matrix. (C) Example receptor activations for two example olfactory signal distributions. Left, low correlation: odor mixtures are generated by 50 relevant odor sources. Right, high correlation: odor mixtures are generated by 5 relevant odor sources. (B) corresponds to right-hand panel of (C). (D) Mutual information as a function of receptor expression levels. Blue lines: sampling. Orange lines: Gaussian approximation. Odorant signal correlation *c*signal = .95, response noise variance *vn*^2^ = .1. Left panel (clean environment) has environmental noise variance 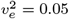, right panel (noisy environment) has environmental noise variance 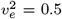 (E) Optimal co-expression (black) vs single-expression (white). Orange line: boundary predicted by the Gaussian approximation.

In our sparse odor model (Fig. 4A), the odorant signal covariance matrix has a low-rank structure with many positively correlated odorant pairs (Fig. 4B, inset). Positive off-diagonal correlations result from pairs of odorants which are emitted by the same odor source. To control the level of stimulus correlation we adjust the number of sources in the signal and the number of odorants emitted by each source. When the olfactory signal consists of fewer sources, each of which emits more odorants, the average correlation between odorants increases.

The affinity of each receptor for each odorant is defined by an affinity matrix **A**, where *a*_*ij*_ describes the affinity of receptor *i* for odorant *j*. Natural receptor-odorant affinities exhibit complex structure that is difficult to reproduce synthetically, so we used empirical data for our receptor affinity matrix **A**. In the absence of data on *Ae. Aegypti* receptor affinities, we used data from another mosquito species, *A. Gambie* sensu lato (s.l.) complex. Specifically, we used receptor affinities from Carey et al. [43], which reports the responses of *m* = 50 receptor types to a battery of *d* = 110 single odorants (Fig. 5B). The matrix entry *a*_*ij*_ reptorts the affinity of receptor *i* for odorant *j* in terms of the change in firing rate from its baseline firing rate. Since the correlation structure of our olfactory stimulus model does not necessarily match the natural correlations of those 110 odorants in the environment, we shuffled the receptor identities to set the entries of our model affinity matrix **A**.

**Fig. 5.**
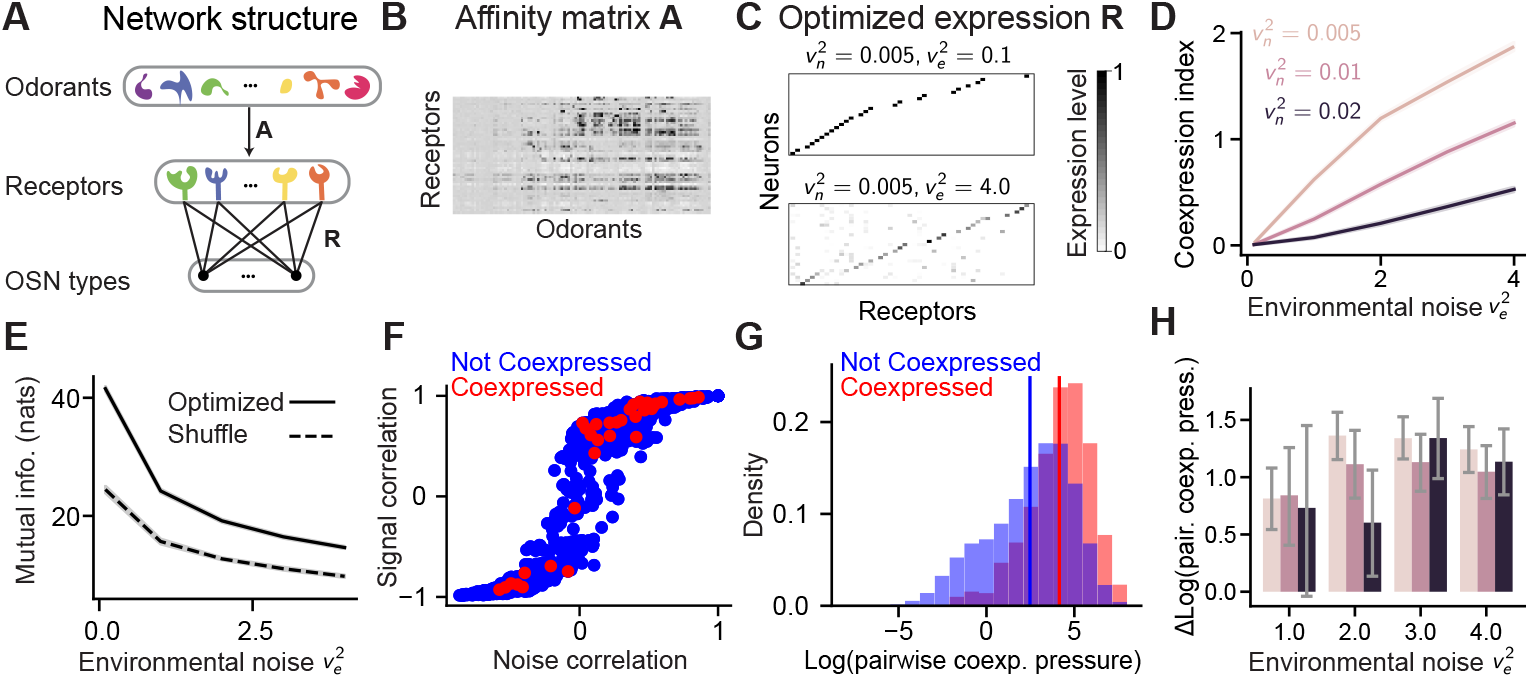
In larger networks, co-expressing pairs or receptors with high signal correlation relative to noise correlation becomes optimal in noisy environments. (A) Network structure. The affinity matrix **A** describes binding affinities for each receptor/odorant pair. Each neuron type has an expression profile described by a row of the expression matrix **R**. (B) Receptor affinity matrix from [43]. (C) Expression matrices resulting from optimizing the mutual information. Top, *ve*^2^ = 0.1, *vn*^2^ = .0205, Bottom, *ve*^2^ = 4.0, *vn*^2^ = .005. (D) Co-expression index (Eq. 33) as a function of environmental noise level *ve*^2^ and neural noise level *vn*^2^. Error bars are 95% confidence intervals. (E) Mutual information from optimized co-expression matrix (solid line) and shuffled co-expression matrix (dashed line) as a function of environmental noise level *ve*2. Shown for neural noise *vn*^2^ = 0.005. Plot shown for *vn*^2^ = 0.01, 0.02 in Fig. S2. (F) Scatter plot of signal correlation **A∑**_*xx*_**A***T* versus noise correlation **AA***T* for all pairs of receptor for one trial. Red: receptor pairs co-expressed in optimized expression matrix **R** for 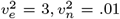. Blue: non-co-expressed pairs. (G) A histogram of the pairwise co-expression pressure for co-expressed pairs (red) and non-co-expressed pairs (blue), pooled over all trials and all values of neural noise and environmental noise. Plotted with the *x* axis on a logarithmic scale. The mean of each distribution is indicated with a vertical line–the mean over co-expressed pairs is 4.03, while the mean over non-co-expressed pairs is 2.33. (H) Difference in the mean value of the log pairwise co-expression pressure for co-expressed, non-co-expressed pairs for all levels of 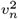 and 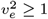. Error bars are 95% confidence intervals. Histograms for all values of 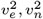 are available in Supplementary Figure S3.

As in the previous section, receptor activations are characterized by their noise correlation (driven by affinity overlap) and signal correlation (driven by both affinity overlap and odorant signal correlation). We compared two olfactory signal distributions with different levels of correlation, which drive different levels of signal correlation between the receptor activations without changing the noise correlation (Fig. 4C).

We investigated the impact of three factors–environmental noise variance *v*_*e*_^2^, neural noise variance *v*_*n*_^2^, and stimulus correlation–on the expression patterns that optimize the OSN response information. We estimated the mutual information, via sampling, between odorant concentrations in the relevant source and neural responses for different receptor expression levels (Methods 3.5). For comparison, we also computed the mutual information of a Gaussian approximation to our sparse odor distribution, where the olfactory signal and noise follow Gaussian distributions with variance and covariance matched to our sparse odor model (Fig. 4D). In both the sparse model and the Gaussian approximation, single expression is optimal when environmental noise is low, but co-expression becomes optimal in the presence of environmental noise (Fig. 4D). The Gaussian approximation captures this transition, although it overestimates the mutual information (Fig. 4D, orange vs blue).

For both the high-correlation and low-correlation olfactory signal distributions, we found the pattern of receptor expression that maximizes the mutual information via sampling. As predicted by our earlier analysis (Equations 3 & 4), higher odorant signal correlation, higher environmental noise, and lower neural noise drive co-expression (Fig. 4E, black: sampling; orange, Gaussian approximation).

### 1.4 Optimal receptor expression patterns in OSN populations

We next determine whether our predictions generalize to larger networks that include multiple types of odorants, receptors, and neurons (Fig. 5A). Additionally, we describe how the optimal co-expression pattern is shaped by the odorant signal distribution and receptor affinities.

Each unit in our model represents an OSN type, or a class of neurons with similar expression profiles that likely project to the same glomerulus. Therefore, we set the number of unit types to be less than the number of receptors. Based on the roughly 2:1 ratio of receptor types to glomeruli in *Ae. aegypti*, we model 25 neuron types, or half the number of receptors. The network’s co-expression pattern is described by an expression matrix **R** (Fig. 5A). Each unit’s response to an olfactory stimulus depends on the receptors it expresses and the affinities of those receptors for each odorant (Fig. 5A,B). We use the same model as in the previous section for the affinity matrix and neural response variability, with response noise variance *v*_*n*_^2^. Also as before, we model the odorant concentrations at the OSN as a combination of the olfactory signal and uncorrelated environmental noise, which has variance *v*_*e*_^2^ (Eq. 1). For numerical tractability, we use the Gaussian approximation of the sparse odor model introduced in the previous section (Fig. 4D,E). See Methods 3.1 for a detailed description of the model.

We first ask whether, in this larger network setting, co-expression is still optimal when the environment is noisy relative to neural responses. For a given stimulus covariance matrix **Σ**_*xx*_, we analyzed the optimal expression matrices from a range of environmental and neural noise levels (Fig. 5C). To see how the levels of environmental and neural noise effect whether co-expression or single expression is optimal across the population, we computed a co-expression index of the optimized expression matrix (Methods, Equation 33). The co-expression index of an expression matrix with *m* receptor types ranges from 0, if each neuron single-expresses one receptor, to *m−* 1, if every neuron expresses all *m* receptors equally. In general, the co-expression index captures the average number of receptors each neuron expresses above one.

As predicted by our earlier calculations for the single neuron, the co-expression index of optimized OSN populations increases with the environmental noise level, but decreases with neural response noise (Fig. 5D). We confirmed this result by mathematically examining the optimal co-expression structures in two limiting cases: when either the environmental or neural response noise is much stronger than the other (Methods 3.6). We proved that single-expression is always optimal when neural response noise dominates (or equivalently, when there is little noise in the olfactory environment). Conversely, when environmental noise dominates, the optimal expression matrix typically features co-expression.

We next ask if and how the structure of co-expression reflects the structure of the stimulus. A shuffled version of the optimal expression matrix performs much worse than the optimized expression matrix (Fig. 5E). Co-expression must thus be somehow structured around the stimulus statistics. We analyzed the co-expression pattern in the optimized expression matrix **R** in terms of the *receptor noise correlation* and the *receptor signal correlation*, which are determined by the affinity matrix **A** and the stimulus covariance matrix **Σ**_*xx*_. The receptor noise correlation is the correlation between the pair of receptors when exposed to independent odorants, i.e., it is the correlation between receptors which results only from overlap in receptor affinity but not from any structure in the olfactory stimulus. On the other hand, the receptor signal correlation is the correlation between receptor activation in response to stimuli taken from the olfactory signal distribution, which results from both overlap in receptor activity and structure in the olfactory signal. Noise correlation and signal correlation are positively related since pairs of receptors with overlapping affinities will be co-activated (Fig. 5F). Co-expressed receptors had higher signal and noise correlations than non-coexpressed receptors (Fig. 5F, blue vs red).

To determine whether and how signal and noise correlations drive co-expression in OSN populations expressing receptors with heterogeneous affinities, we generalized our previous co-expression condition, Eq. 5, to allow for arbitrary receptor affinities (Methods 3.4). In the context of a population of OSNs, this co-expression condition is only approximate since it neglects correlations across neurons’ responses. We used this to define an approximate *pairwise co-expression pressure* for each pair of receptors. The pairwise co-expression pressure is greater than 1 if an isolated neuron should co-express the two receptors, and vice versa. We observed that in the full network, co-expressed receptors have a higher mean pairwise co-expression pressure than non-co-expressed receptors across environmental and neural response noise levels (Fig. 5G,H).

### 1.5 Optimal receptor co-expression with sparse odors and nonlinear responses

To analyze optimal expression patterns in populations of OSNs, we made the simplifying assumptions that odorant concentrations in the relevant source, the environmental noise, and neural response noise all followed Gaussian distributions, and that neural responses are linear. Here, we consider optimal receptor co-expression in populations of OSNs with nonlinear responses and the sparse, non-Gaussian olfactory signal and noise distributions described in Section 1.3.

We use a four-layer encoder-decoder architecture, trained with the objective of decoding odor concentrations in the olfactory signal from olfactory receptor responses (Fig. 6A). Each input to the model is a noisy odorant concentration vector, **y** = **x** + ***ξ*** (Eq. 1). The first layer consists of receptor types (modeled using the shuffled affinity matrix from [43]). The second layer corresponds to OSN types and we interpret the optimized weights of this layer as receptor expression weights. As before, the receptor expression weights are non-negative, the total expression level in each neuron class is constrained, and the OSN units have noisy responses (Methods 3.8).

**Fig. 6.**
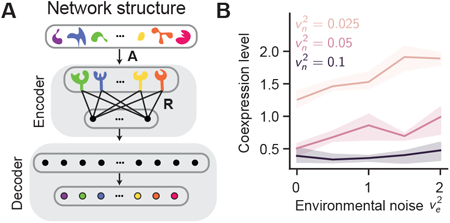
With nonlinear OSN responses, co-expression levels increases with higher environmental noise and lower neural noise. (A) Network architecture: an encoder-decoder structure with two hidden layers, corresponding to the receptors and OSN types. (B) Co-expression level in optimized expression matrix **R** as a function of environmental noise level 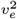 and neural noise level 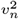.

For varying levels of neural noise *v*_*n*_^2^ and environmental noise *v*_*e*_^2^, we trained the network to estimate the odorant signal concentrations in the input, **x**, and analyze the resulting expression matrix (Fig. S4). As predicted by our earlier analyses, the level of co-expression increases as we decrease the level of neural noise or increase the level of environmental noise (Fig. 6B).

### 1.6 Co-expression of receptors for correlated odorants in flies

Our theory predicts that receptors for odorants that are correlated across behaviorally relevant sources are more likely to be co-expressed. To test this prediction requires 1) a map of odorant-receptor affinities, and 2) measurements of the correlation of those same odorants across relevant natural sources. Dweck et al. published such data for the *Drosophila melanogaster* maxillary palp [44]. Dweck et al. surveyed natural odor sources to identify ligands for all maxillary palp neurons. They found that pB2A neurons respond to furaneol methylether, (-)-camphor, alpha-ionone, beta-ionone, and three unidentified compounds, and that the pB2A response to furaneol methylether is mediated by Or33c while the pB2A response to (-)-camphor, alpha-ionone and beta-ionone is mediated by Or85e [44]. They then measured the presence or absence of 29 odorants across ethologically relevant stimuli for *D. melanogaster*, including 34 fruits extracts, 7 microbe extracts, and 11 fecal extracts.

While the *D. melanogaster* early olfactory system primarily follows the canonical organization, with one receptor type expressed in each neuron, the receptors Or85e and Or33c are co-expressexed in the maxillary palp neuron pB2A (also in *D. pseudoobscura* and *D. simulans* [13]). For each pair of odorants in that dataset, we computed the correlation of the odorant’s presence/absence across all stimuli (Fig. 7A, Methods 3.9). In accordance with our prediction that co-expressed receptors should respond to correlated odorants, the correlation between furaneol methylether, the ligand for Or33c, and (-)-camphor, the main ligand for OR85e, is the largest the largest correlation between pairs of odorants detected by distinct receptors (Fig. 7B).

**Fig. 7.**
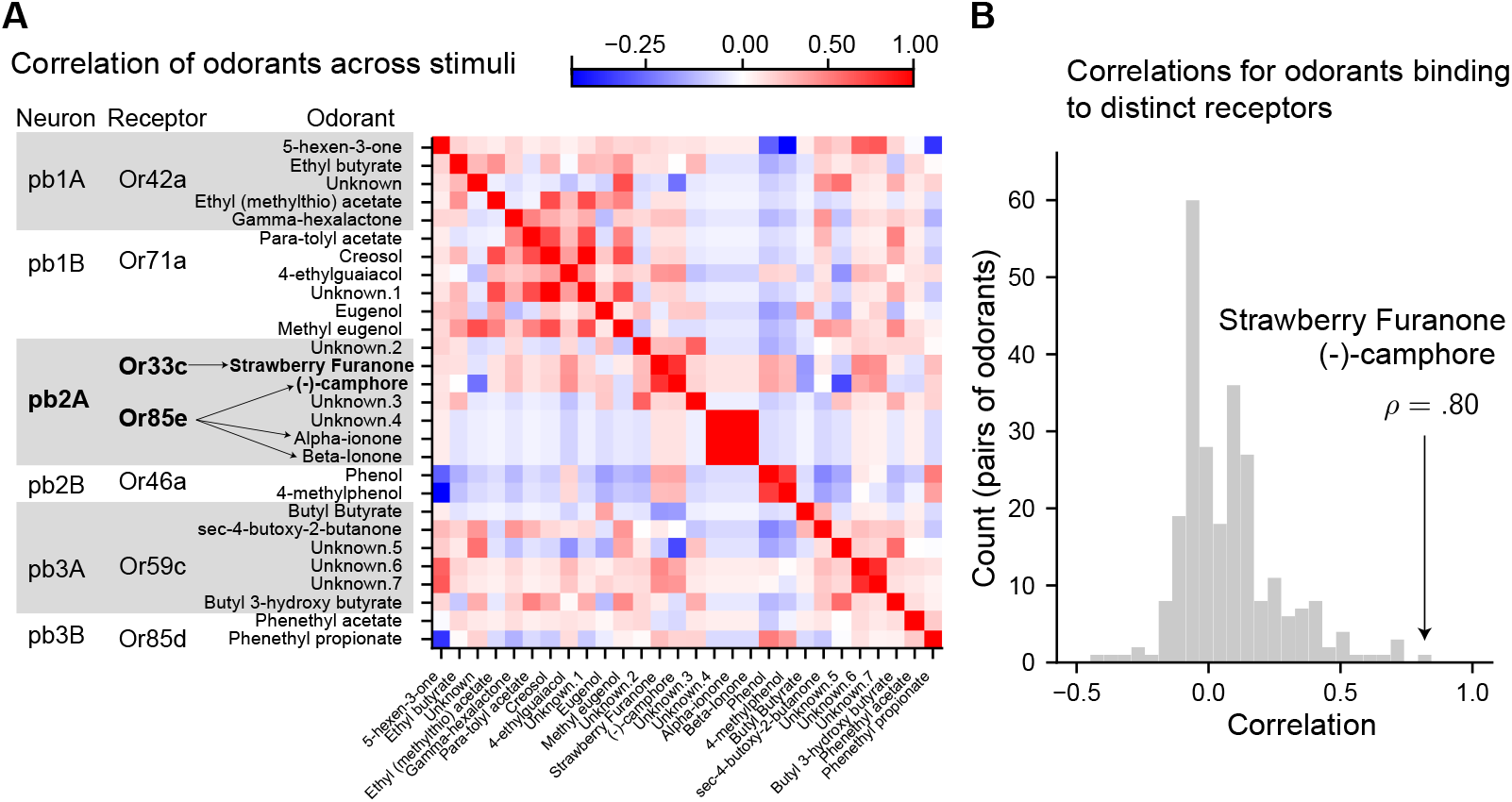
Co-expressed receptors in Drosophila are activated by odorants which are correlated across natural stimuli. (A) Correlation matrix of odorants’ presence/absence across 52 natural stimuli. Odorants are sorted by which maxillary palp neuron they activate. For neuron pb2A, which co-expresses Or33c and Or85e, we plot indicate which odorants bind to each receptor with arrows. Data from Dweck et al 2016 [44]. (B) Histogram of correlations from the matrix in panel (A), restricted to pairs of odorants which bind to the distinct receptors. The correlation between strawberry furanone and (-)-camphore, ligands for Or33c and Or85e respectively, is indicated with an arrow.

### 1.7 A trade-off between stimulus alignment and response gain

We have shown that, in a simple model, co-expression of receptors for correlated odorants is optimal when behaviorally irrelevant fluctuations in odorant concentrations (environmental noise) are the dominant source of noise in OSN odor responses. In contrast, the canonical single-receptor expression motif is optimal when intrinsic variability in neural responses is the dominant noise source. Why are those two expression motifs optimal in those respective scenarios?

We first examine this question for a single OSN type that can express multiple receptors, with weights *r*_1_, …, *r*_*n*_. While a single OSN may respond to many odorants via broad receptor affinity and/or receptor co-expression, the neuron’s firing rate is a one-dimensional variable and cannot directly encode the concentration of every odorant in the olfactory stimulus. To understand how our model OSN deals with this challenge, we decompose their stimulus responses into two stages: 1) a projection into the neuron’s *perceptual subspace* and 2) a scaling by the neuron’s *response gain*.

Different combinations of receptor affinities, expression levels, and odorant concentrations can yield the same net receptor activation (Fig. 8A). Due to this weighting, an OSN’s activity reflects the virtual concentration of an *effective odor blend*, with odorant concentrations weighted by the receptors’ affinities and expression levels. This effective odor blend lies on a surface within the space of odorant concentrations, which we call the OSN’s *perceptual subspace* [45]. The weighting of odorant concentrations by affinities and expression levels projects a stimulus’ odorant concentrations into the OSN’s perceptual subspace (Fig. 8B). Different odorant blends with the same net receptor activation occupy the same position in the perceptual subspace (Fig. 8B).

**Fig. 8.**
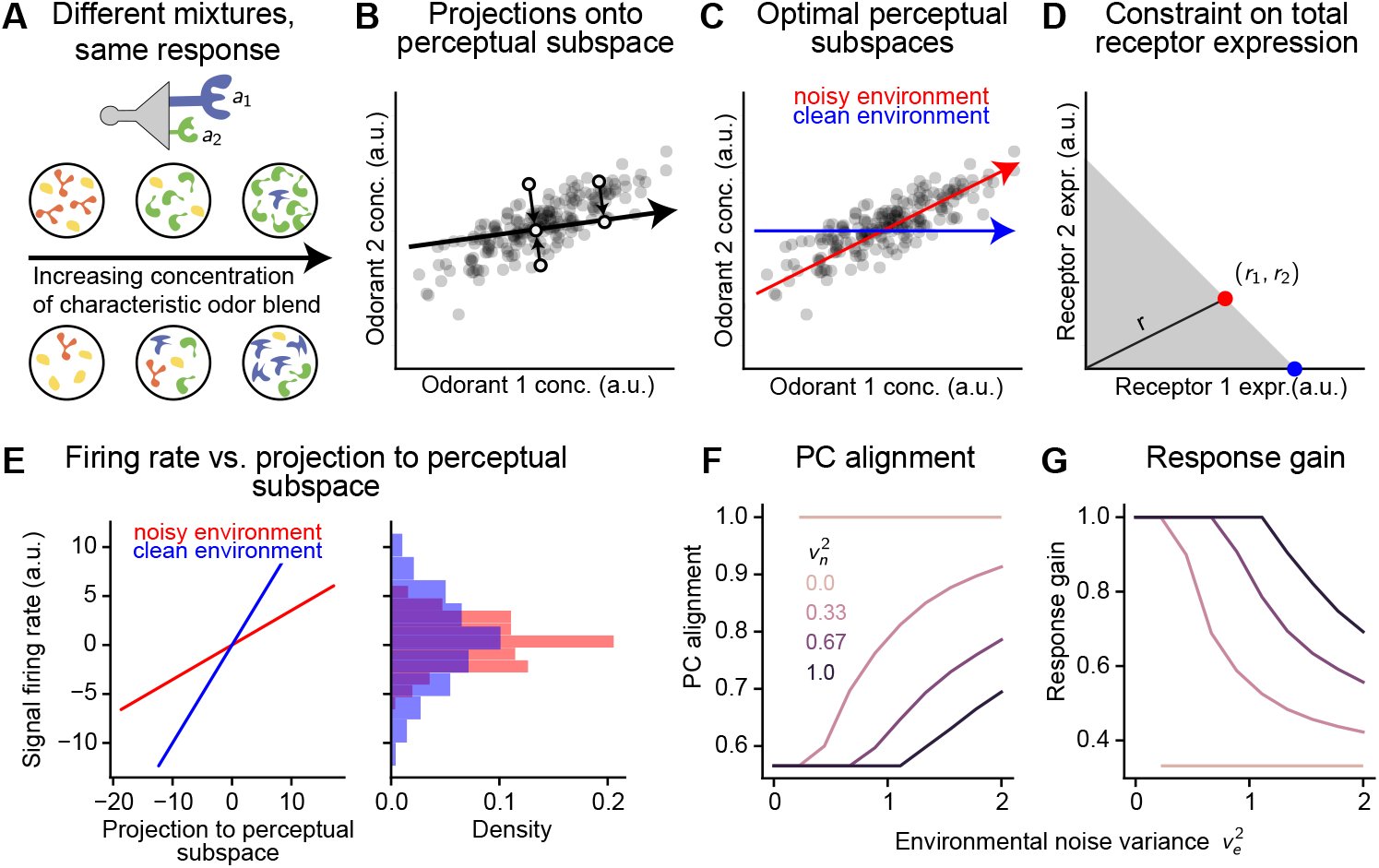
Optimal expression patterns result from a trade-off between maximizing PC alignment maximizing response gain. (A) OSN responses to odor mixtures reflect the concentration of an effective odor blend with odorant concentrations weighted by receptor expression levels and affinities. (B) A single neuron’s response to an odor mixture (shaded points) depends only on its projection (open circles) to the neuron’s perceptual subspace (black line). (C) Perceptual subspaces generated by co-expression vs single-expression. Point cloud: odor concentration vectors from the olfactory signal distribution. Red (blue) line: optimal perceptual subspace for a clean (noisy) environment. (D) Receptor expression weights are limited by the bounded total expression weight (shaded region). Red (blue) dot: optimal expression weights for a noisy (clean) environment. (E) Firing rate vs effective input concentrations (projection to perceptual subspace) for the optimal expression patterns in noisy (red) vs clean (blue). Right: Distribution of OSN rates. (F) The principal component alignment of the optimal expression matrix as a function of environmental noise level and neural noise level. (G) Response gain of the optimal expression matrix as a function of environmental noise level and neural noise level. Optimization as in Fig. 5C-G

We begin by analyzing the optimal perceptual subspace (that corresponding to the optimal receptor expression weights) in the limiting cases with only one noise source, either environmental or response (Methods 3.6). When intrinsic response noise dominates, the optimal OSN expresses one receptor and its perceptual subspace is aligned with the axis corresponding to the highest-variance odorant in the signal (Fig. 8C, blue; Equation 4; Methods 3.6). Why, in this case, is it optimal for the OSN to ignore the other odorants?

The neural response is determined by scaling the effective odorant concentration by the OSN’s *response gain*. Due to the standard mathematical definition of the projection operation, the response gain is the 1 (Fig. 8D, shaded region; illustrated for *n* Euclidean length of the receptor expression vector: 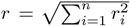. The total receptor expression is constrained via 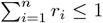 (Fig. 8D, shaded region; illustrated for *n* = 2 receptors). Due to that constraint, the response gain is maximized when the OSN expresses a single receptor (Fig. 8D,E; blue vs red). So, the response gain of the clean environment’s optimal (single-expressing) OSN is higher than the response gain of the noisy environment’s optimal (co-expressing) OSN (Fig. 8E, blue vs red). This causes the distribution of expected neural responses across the stimulus distribution to be more spread out (Fig. 8E, right). In turn, this makes the response more robust to intrinsic neural noise.

When the olfactory environment is the dominant noise source, on the other hand, the optimal perceptual subspace is aligned with the first principal component of the odorant signal **x** (Fig. 8C, red). This provides robustness to the environmental fluctuations in odorant concentrations, since the perceptual subspace is aligned specifically to the dominant axis of behaviorally relevant signals. We measure the PC alignment as cos(*ϕ*), where *ϕ* is the angle between the leading principal component of the olfactory signal distribution and the perceptual subspace. The PC alignment is 1 when the perceptual subspace is perfectly aligned with that leading PC and 0 when the perceptual subspace is orthogonal to the leading PC of the odorant signal. We plot the PC alignment of the optimal perceptual subspace as a function of the environmental noise variance *v*_*e*_^2^ and the neural noise variance *v*_*n*_^2^ for one (linear) neuron with optimized expression matrices (Fig. 8F). We see that the PC alignment of the optimal expression pattern increases with the environmental noise variance and decreases with the neural noise variance. Conversely, we see that the optimal expression pattern’s response gain decreases with the environmental noise variance, while it increases with the neural noise variance (Fig. 8G). This indicates that increasing the response gain provides robustness to neural noise at the cost of decreasing robustness to environmental noise.

Indeed, we can see this trade-off directly in our one-neuron/two-receptor model from Section 1.1, where the odorants have signal correlation *c >* 0. In that case, the constraint on total receptor expression levels implies that

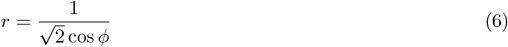

where *r* is the response gain and cos *ϕ* the principal component alignment.

Unless the first principal component of the olfactory stimulus distribution is aligned with one odorant axis, i.e., one odorant is independent of all others, a neuron whose perceptual subspace is aligned with the leading principal component exhibits co-expression. To align with the signal’s first principal component, the receptor expression vector generally needs to sample multiple axes in odorant space. This can occur through broad receptor affinities and/or co-expression. Due to the constant on total expression levels, co-expression forces the expression vector to be shorter than that of a single-expressing neuron (Fig. 8D, red vs blue). The length of the expression vector is exactly the response gain described above. Thus, co-expression is optimal when environmental noise is high relative to neural noise because it enables PC alignment, while single expression is optimal when environmental noise is low relative to neural noise because it maximizes response gain. These results generalize directly to the average response gain and population perceptual subspace of OSNs and receptors with abritrary sensing matrices (Section 3.6; Fig. S5). We speculate that similarly, broadening receptor affinities may come at the cost of decreasing the amplitude of responses to single odorants.

In summary: under a constraint on total receptor expression levels, the olfactory system must trade off between aligning neural responses to the leading principal components of the stimulus distribution and between maximizing the stimulus-driven variance of neural responses. As we increase the level of environmental noise *v*_*e*_^2^ relative to neural noise *v*_*n*_^2^, PC-alignment is favored over response gain. Because co-expression is typically required for PC-alignment, while single expression maximizes response gain, noisier environments promote co-expression while clean environments promote single expression.

## 2 Discussion

Our results demonstrate that co-expression of olfactory receptors contributes to robustness to noise in the environment, at the cost of decreasing robustness to neural noise. Thus, co-expression should be favored when an organism faces a noisy environment and has invested in reliable neural responses. Further, this co-expression must be structured around natural stimulus statistics, with receptors for correlated odorants more likely to be co-expressed. In this case, the early olfactory system to perform PCA-like dimension reduction on its input.

### 2.1 Predictions

In addition to predicting that co-expressed receptors should correspond to odorants which are correlated across the olfactory signal, our model predicts that co-expression is promoted when the level of environmental noise is high relative to the level of neural noise. Does this help explain which species and subsystems exhibit co-expression and which exhibit single expression?

Systems which exhibit co-expression may be specialists, rather than generalists. For instance, *D. melanogaster*, which primarily exhibit single expression, feed on a variety of rotting fruit and decaying plant matter, as well as on some animal sources of food [46, 47]. On the other hand, female *Ae. aegypti* mosquitoes are specialists, primarily feeding on human blood. Our model does not explicitly incorporate innate valence of odorants or odor mixtures. However, the division of the olfactory environment into signal and noise will differ between specialists and generalists. For specialists, fewer types of odor sources are ethologically relevant. More irrelevant odor sources will thus contribute to olfactory noise. Thus, we expect specialist species to experience higher levels of environmental noise and higher correlations between relevant odorants, promoting co-expression.

For example, mosquito species vary in the extent to which they specialize on a single host species for blood feeding, and only female mosquitoes blood-feed. Thus, the prediction that specialization, and therefore a noisier environment, promotes co-expression could be tested by examining whether or not co-expression also appears in mosquito species which feed on a wider variety of organisms.

Changes in the neural noise level can also drive changes in whether co-expression is optimal. In adult *D. melanogaster*, the OSN ab10B co-expresses the receptors Or49a and Or85e, both of which bind to ligands emitted by parasitoid wasps of the genus *Leptopilina* [14]. On the other hand, in *D. melanogaster* larvae, the corresponding OSN single-expresses Or49a. Because the adult flies have 2400 OSNs and larvae have only 21 [48], adult flies can pool the responses over a larger population of OSNs. Because pooling responses of many OSNs can average out intrinsic fluctuations in individual neurons’ activity, adult flies should effectively have a lower level of neural response noise relative to fly larvae. This is consistent with our prediction that co-expression is favored by lower levels of neural noise relative to environmental noise.

In Section 1.1, we note that when odorants do not have equal variance, expression is biased towards expressing more of the receptor for the higher-variance odorant. This mirrors results from [49], which predicts that the abundances of olfactory receptors in mammals should adapt to express more OSNs with receptors for higher-variance odorants when there is more neural noise. In *Ae. aegypti* OSNs, overall expression levels vary between receptors [22]. We expect that the ligands for more highly-expressed receptors have higher variance in the relevant olfactory environment.

### 2.2 Can efficient coding explain other violations of the canonical organization?

In addition to violating the “one-neuron-one-receptor” principle, many species violate the “one-neuron-one glomerulus” aspect of the canonical organization. In locusts (*Schistocerca gregaria*), OSNs project to multiple glomeruli, and projection neurons innervate multiple glomeruli [50, 51]. Divergent projection of OSNs to multiple glomeruli is also the norm in many amphibian species [52–54], and is observed in the accessory olfactory bulb in rodents [55]. In rodents, convergence of OSNs expressing distinct olfactory receptors is observed in the necklace glomeruli within the main olfactory bulb [19]. Here, we have used information theory to investigate how the relative levels of environmental noise and neural noise favor single-expression versus co-expression, motivated by the widespread co-expression found in *Ae. aegypti*. A simple generalization of our model would be to view each OSN type instead as a glomerulus. Similar principles from information theory may be useful in understanding why in some systems, glomeruli pool inputs from multiple OSN types.

### 2.3 Randomness versus structure

Many models of the olfactory system show that random “mixed-selectivity” performs well in stimulus classification. For instance, the connectivity between projection neurons in the antennal lobe and Kenyon cells in the mushroom body is often described as sparse and random. This sparse, random connectivity has been found to maximize dimensionality of the Kenyon cell layer, which in turn makes it easier for downstream neurons to perform associative learning based on Kenyon cell output [56]. Additionally, this sparse, random structure makes it possible to decode odor identity from Kenyon cell activity [57]. Similarly, when olfactory signals are sparse, the diffuse binding of olfactory receptors to multiple odorants makes it possible to decode olfactory input from receptor activation, even when the number of receptor types is small relative to the number of odorants due to the principle of compressed sensing [58, 59].

In contrast, but similarly to [60], we find that the co-expression pattern must be structured around the statistics of natural stimuli, and that random co-expression does not lead to a good OSN code. Recent work supports the idea that other parts of the olfactory system are structured around the statistics of natural stimuli, rather than purely random or structured around chemical similarities [61, 62]. In particular, a measure of metabolic similarity predicts receptor affinity better than chemical descriptors [62]. If molecules with a high level of metabolic similarity are more likely to co-occur in natural stimuli, then this structure to receptor affinities may perform better in noisy environments.

### 2.4 Limitations

In this study, we made several simplifying assumptions. First, we fixed the number of OSN types. Co-expression emerges as optimal only when the number of OSN types is fixed to be smaller than the number of receptors. However, the number of OSN types and olfactory glomeruli varies across insect species. Thus, we have not explained why in *Ae. aegypti*, the number of OSN types or glomeruli did not increase in order to accommodate a larger number of olfactory receptors. While increasing the number of OSN types and glomeruli to match the number of receptors should increase the mutual information about the stimulus, there are also benefits to having a bottleneck. Bottleneck architectures de-noise and de-correlate input to the Kenyon cells while decreasing the number of synapses required [60]. Thus, the informational benefit derived from having a larger number of distinct OSN types should be weighed against the costs of having more glomeruli.

Second, our model neglects the temporal dynamics of odor concentrations and of OSN activity. Animal’s ability to track temporal structure in odor concentrations underlies their ability to navigate turbulent odor plumes [63–66] and can be used to perform olfactory source separation [67, 68]. Likewise, many proposed mechanisms for coding for odor identity, such as the primacy coding model, depend on temporal pattern of activation in the OSNs and glomeruli [69, 70]. By determining optimal responses to static odor concentrations, our theory describes equilibrium coding for concentration fluctuations in linear regimes. OSNs are spiking neurons, and it is possible that the discrete nature of spiking fluctuations impacts their odor coding, especially in low-rate regimes (where the Gaussian approximation we employed is a poor one). A more detailed analysis of noise distributions may reveal that details of the noise structure determine whether or not co-expression is optimal.

Finally, in several sections we focused on maximizing mutual information under linear odorant-receptor-neuron models. Odorant-receptor interactions are nonlinear due to, for example, competitive binding [71, 72], which may also shape optimal receptor expression patterns. An intriguing hypothesis is that those interactions are also adapted to the natural statistics of relevant stimuli [73, 74].

## Acknowledgements

C.L. is supported by the Swartz Foundation and the BU Center for Systems Neuroscience. M.A.Y. is supported by the Searle Scholars Program, the Richard and Susan Smith Family Foundation, the Esther A. & Joseph Klingenstein Fund, the Simons Foundation, the Alfred P. Sloan Foundation, and the Pew Charitable Trusts. We thank Tyler Hill, Jack Giblin, and Wesley Alford for helpful feedback.

## Code availability

Software and data are available at https://github.com/lienkaemper/mosquito MI.

## 3 Methods

### 3.1 Model details and mutual information

Let *d* be the number of odorants, *m* the number of receptors, and *n* the number of neurons. We model the olfactory signal as an odorant concentration vector **x** ∈ ℝ^*d*^ sampled from a multivariate normal distribution *𝒩* (***µ***_*x*_, **Σ**_*xx*_). Independent Gaussian noise is added so that the odor concentrations which reach the neuron are

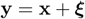

with 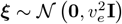.

Each neuron’s response to a stimulus, i.e. to an odorant concentration vector, is linear in the odorant concentrations, weighted by both the receptor expression levels and the receptor affinities. Additionally, each neural response is corrupted by independent Gaussian noise with variance *v*_*n*_^2^. The expression levels of each receptor are described by a *n × m expression matrix* **R**, with *r*_*ij*_ giving the expression weight of the *j*-th receptor in the *i*-th neuron. We require all receptor expression weights to be non-negative, and constrain the total expression level in each neuron, i.e.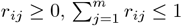. The affinity of each receptor for each odorant is described by the *m × d affinity matrix* **A**, with *a*_*ij*_ giving the affinity of receptor *i* for odorant *j*. Thus, the response of neuron *i* to the received odorant concentration vector **y** is

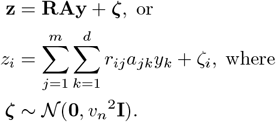

**Table 1.** Model variables and key terms.

| Term | Variable |
| --- | --- |
| Odorant signal | $\mathbf{x}$ |
| Odorant noise | $\xi$ |
| Olfactory input | $\mathbf{y}$ |
| Neural response | $\mathbf{z}$ |
| Neural noise | $\zeta$ |
| Receptor expression | $\mathbf{R}, r_{ij}$ |
| Receptor affinity | $\mathbf{A}$ |
| Stimulus covariance | $\Sigma_{\mathbf{x}\mathbf{x}}$ |
| Environmental noise variance | $v_e^2$ |
| Neural noise variance | $v_n^2$ |
| Correlation between odorants | $c$ |
| Number of neurons | $n$ |
| Number of odorants | $d$ |
| Number of receptors | $m$ |
| Odorant stim. variance (unequal case) | $\sigma_1^2, \sigma_2^2$ |
| Sigmoid function | Sigmoid |
| Probability | $p$ |
| Mutual information | $I_R(\mathbf{x}, \mathbf{z})$ |
| Affinity overlap | $a$ |
| Response gain | $r$ |
| Coexpression level | co-expression level |

We quantify the information that the neural responses **z** contain about the stimulus **x** via the mutual information

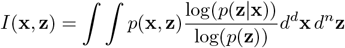

Because both noise sources are Gaussian and the neurons are linear, the joint distribution of the odor concentrations and firing rates is also a Gaussian distribution, with covariance matrix

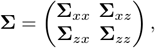

where

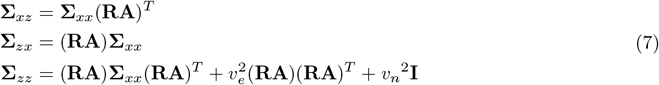

Here, (**RA**)**Σ**_*xx*_(**RA**)^*T*^ is the covariance matrix of the mean responses: the response signal covariance. The response noise covariance is 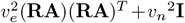 For Gaussian distributions **x** and **y** with joint covariance matrix **Σ**, the mutual information *I*(*x, y*) satisfies

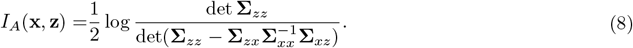

Note that the mutual information does not depend on the mean input or mean response. For one dimensional Gaussian distributions, the mutual information relates to the signal to noise ratio as 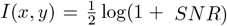 This is generalized to higher dimensional Gaussian distributions by the identity

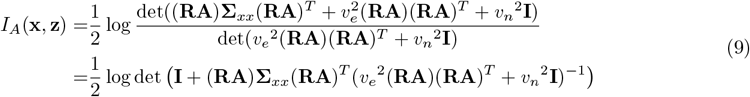

While we constrain the total amount of expression in each neuron via 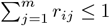, the mutual information is non-decreasing under a scaling all receptor affinities within a neuron by a constant factor *c* ≥ 1. Thus, the optimal mutual information is always attained on the boundary of this region, i.e., where 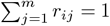

### 3.2 Optimal expression for one neuron with two receptors

For the olfactory signal, we first take two odorants with unit variance and correlation *c* across natural stimuli, i.e., 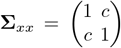. We consider a one-to-one relationship between odorants and receptors, i.e. **A** = **I**. Because we only consider one neuron in this example, we drop the first index, writing our expression as **R** = (*r*_1_, *r*_2_). By Equation 9, the mutual information is

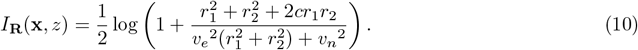

To find the optimal expression (*r*_1_, *r*_2_) as a function of *c, v*_*e*_^2^, *v*_*n*_^2^, we make the substitution 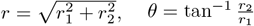. Then

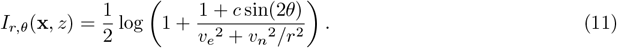

Now, due to the constraint *r*_1_ + *r*_2_ = 1, we have *r* (cos *θ* + sin*θ*) = 1. Then 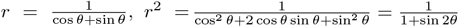. Substituting this in to equation 11, we can eliminate *r*:

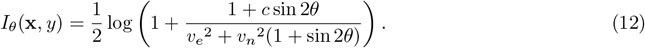

Now, since 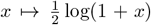 is a monotone function, maxima of *I*_*θ*_(**x**, *z*) correspond to maxima of the function

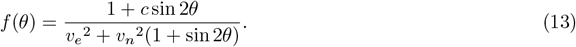

Critical points, including maxima, occur at zeros of:

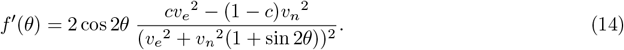

Since 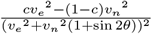 has no zeros unless *cv*_*e*_^2^ − (1 − *c*)*v*_*n*_ ^2^ = 0, all zeros of *f′* coincide with zeros of sin 2*θ*. When *cv*_*e*_^2^ − (1− *c*)*v*_*n*_^2^ *>* 0, the only local maximum of *f ′* occurs at the local maximum of sin 2*θ*, i.e. at *θ* = 45^*°*^. When *cv*_*e*_^2^ − (1 −*c*)*v*_*n*_^2^ *<* 0, there are two local maxima of *f ′* coinciding with the two local minima of sin 2*θ*, i.e. at *θ* = 0^*°*^, 90^*°*^. When *cv*_*e*_^2^ − (1 −*c*)*v*_*n*_^2^ = 0, the mutual information is constant for all *θ*. This results in our co-expression condition, Equation 3.

### 3.3 Optimal expression for one neuron: unequal odorant variances

Next, we consider how this result changes when the two odorants do not necessarily have the same variance. In general, the stimulus covariance matrix has the form

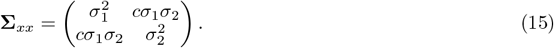

Without loss of generality, assume 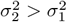. By Equation 8, we have

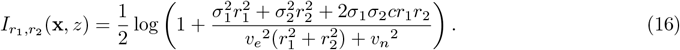

Using the constraint *r*_1_ + *r*_2_ = 1, we eliminate *r*_2_ to write 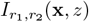 solely in terms of *r*_1_. Again, since 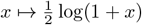 is monotone, maxima of *I*_*r1, r2*_ (**x**, *z*) occur at maxima of

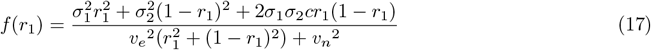

To find the expression pattern *r*_1_, *r*_2_ where mutual information is maximized, we find the zeros of *f′*. Since 0 ≤ *r*_1_ ≤ 1, some level of co-expression is optimal whenever *f ′*has a root in (0, 1) where *f″ >* 0. We have

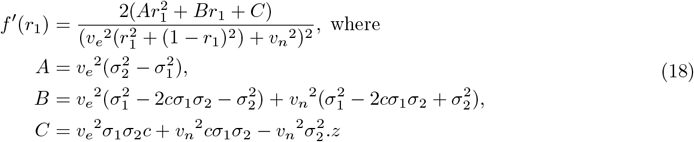

Since the numerator of *f′* is quadratic in *r*_1_, we can find the roots using the quadratic formula. Since we assumed that 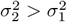, the smaller root of *f*^*′*^is a local maximum, while the larger root is a local minimum. To find an analogue of our co-expression condition (Equation 3), we solve for the values of 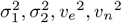, and *c* so that *f′* has a zero at the origin, i.e. the point where the maximum moves out of the region (0, 1). Since the zeros of *f′* are at 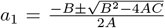, there is a root at zero when 4*AC* = 0. Since *A ≠* 0, this requires *C* = 0. Setting *C* = 0, we obtain the co-expression condition

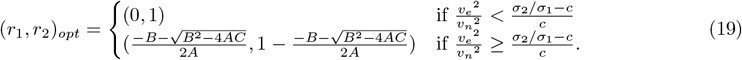

When *v*_*n*_^2^ = 0, co-expression is always optimal, and (*r*_1_, *r*_2_)_*opt*_ is the leading eigenvector of **Σ**_*xx*_. This means that the neural activity *y*(**x**) is the alignment of **x** with the first principal component the inputs.

### 3.4 Optimal expression for one neuron: overlapping receptor affinities

Now, we relax the assumption that there is a one-to-one relationship between odorants and receptors. We begin with an affinity matrix

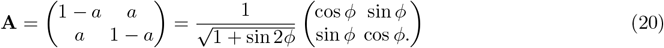

and later consider general affinity matrices. This form for the affinity matrix, in which the row sum does not change as *a* is varied, ensures consistent behavior when both the two odorants and the two receptors are identical. Here, the overlap angle *ϕ* describes each receptor’s sensitivity to the other receptor’s odorant, i.e. when *ϕ* = 0^*°*^, receptor one is a perfect detector for odorant one, and when *ϕ* = 45^*°*^, both receptor are equally sensitive to both odorants. As in our first example, we use the stimulus covariance matrix 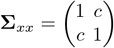.

As before, we work in polar coordinates, setting 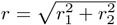 and 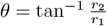 We calculate the mutual information via Equation 9 as

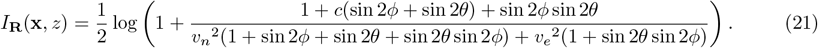

Again, we set

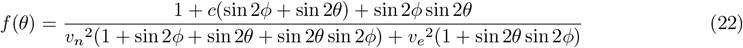

and compute the zeros of

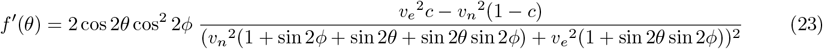

Thus, we obtain the same co-expression condition as we did without the sensing matrix. However, increasing the overlap angle *ϕ* between the receptors decreases the absolute value of *f′* (*θ*). Thus means that as the overlap between receptors becomes greater, the difference in mutual information between co-expression and single expression becomes less.

When there are more than two odorants, the receptor affinity matrix has the more general form

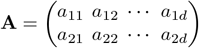

where *d* is the number of odorants. In this case, we can describe whether or not co-expression is optimal in terms of the *signal correlation* and the *noise correlation* between receptors. We compute the signal and noise covariance matrices of the receptor activations as

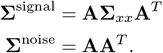

We then normalize these to obtain values for signal and noise correlation:

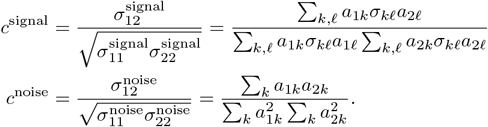

Notice that the noise correlation *c*^noise^ is driven purely by tuning overlap. On the other hand, we can decompose numerator of the signal correlation in to two parts, one due to correlation in the signal and one due to the tuning overlap:

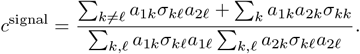

The term ∑_*k ≠ 𝓁*_ *a*_1*k*_*σ*_*k 𝓁*_*a*_2 *𝓁*_ is driven by correlations in the signal: higher correlations between relevant odorants, i.e. higher values of *σ*_*kl*_, that drive both receptors increase signal correlation. On the other hand, the term ∑_*k*_ *a*_1*k*_*a*_2 *k*_*σ*_*kk*_ is determined by the way that the tuning overlap relates to the variance of odorants across the signal:_*∑k*_ *a*_1*k*_*a*_2*k*_*σ*_*kk*_ is enhanced relative to the noise covariance ∑_*k, 𝓁*_ *a*_1*k*_*σ*_*k 𝓁*_*a*_2 *𝓁*_ if the receptors overlap on high-variance odorants, while it is decreased relative to the noise covariance if the receptors overlap on low-variance odorants.

The mutual information is a function of the signal and noise covariance matrices, **Σ**^signal^ and **Σ**^noise^, the environmental and neural noise variances 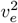 and 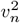, and the receptor expression weights *r*_1_ and *r*_2_:

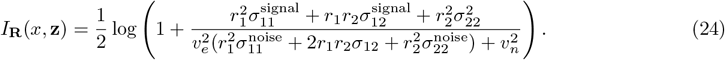

Using similar techniques as above, we find conditions for when single-expressing receptor 1 is optimal, when single-expressing receptor 2 is optimal, and when co-expression is optimal. By evaluating at **R** = [1, 0] and **R** = [0, 1], we determine that single expression of receptor 1 outperforms single expression of receptor 2 when

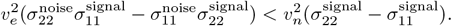

To determine when co-expression is optimal, we set *r*_2_ = 1 − *r*_1_ and

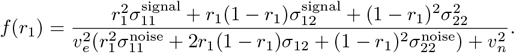

As in section 3.3, we determine whether mutual information can be increased from the optimum achievable with single expression by determining the sign of *f*′ (*r*_1_) at *r*_1_ = 1 if single-expressing receptor 1 is optimal and at *r*_1_ = 0 if single expressing receptor 2 is optimal.

We have that co-expression is better than single-expressing receptor 1 (*f*′ (1) *<* 0) when

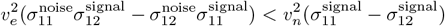

and co-expression is better than single-expressing receptor 2 (*f*′ (0) *>* 0) when

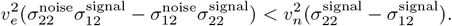

In the main text we focus on the case where both receptors have equal signal variance 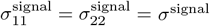 and equal noise variance 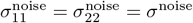. In this case, co-expression is optimal when

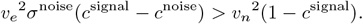

Thus, signal correlation must always be greater than noise correlation in order for co-expression to be optimal. The earlier result for the affinity matrix 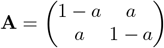 is a special case of this calculation.

### 3.5 Estimating mutual information via sampling

Our olfactory signal and noise distributions are each defined defined by a set of odor sources, each with a characteristic odor blend which is sparse in the set of odorants We first generate a sparse *d × n*_*sources*_ matrix **T**. Each column of **T** represents an ethologically relevant odor source, while each row of **T** represents an odorant. Each source has a characteristic odor blend which containing a small number of odors. To model this, we sample each entry of **T** to be non-zero with probability *p*_*source*_. We set the non-zero entries with a uniformly distributed values in the range [0, *c*_*max*_], *c*_*max*_ = 10.

To sample from the olfactory signal distribution, we first add Gaussian noise with variance *κ*^2^ to the positive odor concentrations in each source, and then ensure positive odor concentrations in the noisy sources by setting all negative concentrations to zero. We then take a sparse linear combination of noisy versions of the signal odor sources, with each odor source participating in the sample with a probability of *p*_*sample*_. Each source which participates in the sample has a weight *k*_*i*_ drawn uniformly from [0, 1]. A sample is then given as **x** = *T* **k**. Likewise, we sample from the olfactory noise distribution by taking a sparse linear combination of the noise odor sources. We compute the covariance matrix of the olfactory signal, and denote it as **Σ**. We scale the olfactory noise by an overall factor 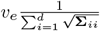, so that the average variance across odorants is *v*_*e*_^2^.

We estimate the mutual information between olfactory input and neural response via the formula

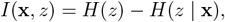

where *H*(**z**) is the entropy of the neural responses, i.e. *H*(**z**) = ∫*p*(*z*) log(*p*(*z*))*dz*, and *H*(*z* | **x**) is the entropy of the neural responses conditional on the olfactory signal, i.e. *H*(*z* | **x**) = ∫*p*(**x**) *∫p*(*z*|**x**) log(*p*(*z*|**x**))*dzd***x**. We estimate the value of both of these integrals via sampling, noting that *H*(*z*) = ⟨log(*p*(*z*))⟩_*x*_ and *H*(*z* | **x**) = ⟨⟨log(*p*(*z*| **x**))⟩_*z*_⟩_**x**_.

Using our definition of the neural response as *z* = **RA**(**x** + *ξ*) + *ζ*, we can estimate *H*(*z*) and *H*(*z* **x**) as a function of the expression pattern **R** by summing over samples of **x**, *ξ*, and *ζ*:

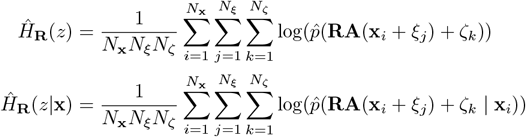

We compute 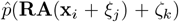 by binning over all samples of **x**, *ξ, ζ* and counting the number of samples in each bin. We compute 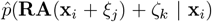 by binning values of **RA**(**x**_*i*_ + *ξ*_*j*_) + *ζ*_*k*_ for all samples of *ξ* and *ζ* for each fixed value of **x**_*i*_.

To approximate the optimal expression pattern, we estimate the mutual information as 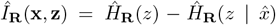 for ten evenly spaced values of *a*_1_ between *r*_1_ = 0 and *r*_1_ = 1. As before, we fix *r*_2_ = 1 − *r*_1_. We say that single-expression is optimal is the maximum value of 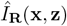 is obtained at either **R** = [1, 0] of **R** = [0, 1], and co-expression is optimal otherwise.

For the low-correlation signal, we used *n*_source_ = 100, *p*_source_ = .05, and *p*_sample_ = .1. For the high-correlation signal, we used *n*_source_ = 5, *p*_source_ = .2, and *p*_sample_ = .4. In both cases, olfactory noise was sampled with parameters *n*_source_ = 1000, *p*_source_ = .05, and *p*_sample_ = .1.

### 3.6 Larger networks: derivations in extremal cases

We now consider networks with more neurons and more receptors. In most cases, we are no longer able to find the optimal expression pattern analytically. The exception is the two extreme cases when one noise source completely dominates the the other, i.e. *v*_*e*_^2^ *≫v*_*n*_^2^ or *v*_*n*_^2^*≫ v*_*e*_^2^. In all these cases, we assume that the number of neuron types *n* is less than the number of receptor types *m*.

#### 3.6.1 Noisy environment

We first consider the case where environmental noise dominates neural noise. Setting *v*_*n*_^2^ = 0 in Equation 9, we have

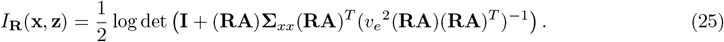

Let **RA** have singular value decomposition **RA** = **UEV**^*T*^, and note that since *m > n*, **U** and **E** are invertible, square matrices while **V** is a rectangular *n × m* matrix. Substituting this in to our formula for mutual information and noting that *U* and *V* are orthogonal matrices, so that **U**^*T*^ **U** = **I** and **V**^*T*^ **V** = 1, we have

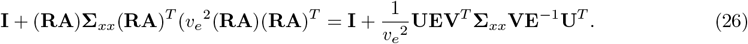

Again using the fact that **U**^*T*^ **U** = **I** and **E** is invertible, we can bring **UE** out as

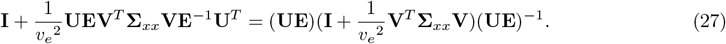

Now, since the determinant is multiplicative, we have

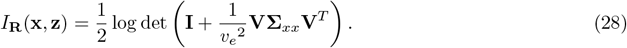

That is, the only way in which the neural affinity matrix **RA** affects the mutual information via its kernel, i.e., which directions in odor space the neural population records and which it discards.

Now, let *λ*_1_ *> > λ*_*O*_ be the eigenvalues of **Σ**_*xx*_, and 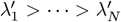 be the eigenvalues of **VΣ**_*xx*_**V**^*T*^. Notice that

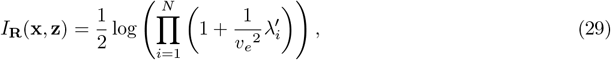

which is an increasing function of all eigenvalues 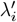. By Cauchy’s interlacing theorem, 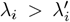 for all *i* = 1, …, *n*. This maximum is achieved when the rows of **V** are the *n* leading eigenvectors of **Σ**_*xx*_.

When 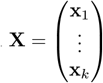 is a set of olfactory stimuli drawn from *𝒩* (**0, Σ**_*xx*_), in the limit of large data *k* → ∞the empirical covariance matrix **X**^*T*^ **X** approaches **Σ**_*xx*_. Then the optimal co-expression matrix **A** satisfies 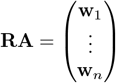 where **w**_1_, …, **w**_*n*_ are the weights of the first *n* principal components of the olfactory stimuli.

Typically, this will not be achievable with single-expression.

Thus, in the limit where environmental noise *v*_*e*_^2^ dominates neural noise *v*_*n*_^2^, co-expression is optimal because it makes it possible for the early olfactory system to perform PCA on its input. One complication is that it is not always possible to achieve this with the constraint that receptor expression levels are positive.

#### 3.6.2 Clean environment

Next, we consider the case where neural noise dominates environmental noise. Setting *v*_*e*_^2^ = 0 in Equation 9, we have

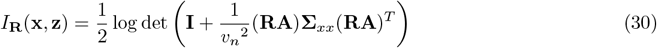

We prove that the optimal expression matrix **R** always at most one nonzero entry per row; that it expresses a single receptor.

##### Proposition 1.

If **Σ**_*xx*_ is full rank and *v*_*e*_^2^ = 0, any expression matrix **R** which maximizes *I*_**R**_(**x, z**) has at most one nonzero entry per row.

*Proof* Suppose to the contrary that there is a matrix **R** which maximizes *I*_**R**_(**x, z**) and which does not have single expression. Let **z \**_*i*_ = (*z*_1_, …, *z*_*i*−1_, *z*_*i*+1_, …, *z*_*n*_) be the activity of all neurons other than neuron *i*. Then by the mutual information chain rule,

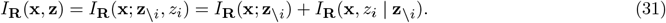

Notice that only the value of *I*_**R**_(**x**, *z*_*i*_|**z**_*\i*_) depends on the expression weights in the neuron *i*. Let Σ_*x*|*z*_ be the covariance matrix of the conditional distribution for *p*(**x**|**z**_*\i*_) and **R**_*i*_ the *i*-th row of **R**. Then

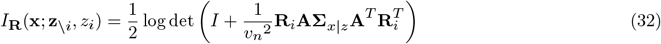

Notice that the signal to noise ratio 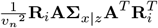 is a convex function of **R**_*i*_. Thus, it is maximized on a vertex of the unit simplex 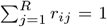, i.e., at single expression. Therefore, it is possible to increase the mutual information *I*_**R**_(**x, z**) by changing one neuron to have single expression, contradicting our assumption.

### 3.7 Larger networks: optimization techniques and analysis

When both *v*_*e*_^2^, *v*_*n*_^2^ *>* 0, we can no longer compute the optimal expression pattern analytically. Instead, we fix a signal covariance matrix **Σ**_*xx*_ and a sensing matrix **A** and numerically find an expression matrix **R** which is a local maximum for the mutual information.

We generate a low-rank signal covariance matrix **Σ**_*xx*_ by computing the covariance across odorants sampled from the sparse mixture distribution as described in Section 3.5, with *c*_*max*_ = 10, *p*_*source*_ =.15, *p*_*sample*_ = 1, and *n*_*source*_ = 25. Notice that in this case, **TT**^*T*^ is a low-rank matrix and its *i, j* element gives the covariance between odorant *i* and odorant *j* over odor concentration vectors **x** sampled i.i.d. as mixtures of the odor sources given by the columns of **T**. To represent the fact that different samples from the same odor source may emit slightly different odor blends, we add a diagonal term to the signal covariance matrix, i.e. **Σ**_*xx*_ = **TT**^*T*^ + *κ*^2^**I**. This is a Gaussian approximation to the sparse odor distribution model which we use in Section 3.5. To produce Fig. 5, we use *d* = 110, *n*_*s*_ = 25, *p*_*source*_ = .15, *c*_*max*_ = 10, *κ* = 0.5. We optimize the mutual information using projected gradient ascent constrained to the simplex 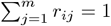 implemented with PyTorch. All our analysis is the average of 50 trials, with different values of the covariance matrix. Learning curves are plotted in Supplementary Figure S1.

We quantify the level of co-expression exhibited by the optimized expression matrix **R** via

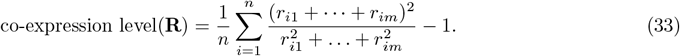

Notice that if each neuron expresses *k* receptors in equal amount, then

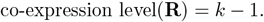

To see how natural stimulus statistics shape the optimal expression pattern, we define the *pairwise co-expression pressure* in terms of the signal and noise covariance of each pair of receptors, as well as the levels of environmental and neural noise. We then compare the pairwise co-expression pressure between pairs of neurons which are co-expressed and pairs of neurons which are not co-expressed. We denote the entries of the receptor-by-receptor signal covariance matrix **Σ**^signal^ = **AΣA**^*T*^ and noise covariance matrix **Σ**^noise^ = *AA*^*T*^ as 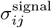 and 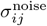 respectively. In Section 3.4, we showed that in a setting with only two receptors and one neuron present, co-expression outperforms single expression of either receptor when

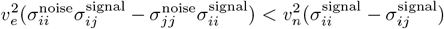

and

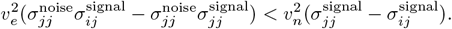

Motivated by this, we define the pairwise co-expression pressure as

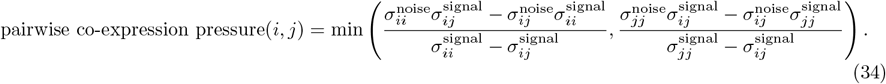

We compare the pairwise co-expression pressure between pairs of receptors that are co-expressed, and those that are not. Because the pairwise co-expression pressure is a ratio, we always plot it on a log scale. Finally, we compare the dimension reduction learned by the neurons via the optimal co-expression matrix **R** to principal component analysis. To do this we compare the perceptual subspace, defined as the row-space of the matrix **RA**, which is a *n* dimensional subspace of odor space, to the principal subspace, defined as the span of the *n* leading eigenvectors of **Σ**_*xx*_. We define the principal component alignment as 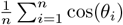, where *θ*_1_, …, *θ*_*n*_ are the principal angles between the principal subspace and the perceptual subspace. The principal component alignment ranges from 0, when the perceptual subspace is orthogonal to the principal subspace, to 1, then the perceptual subspace is identical to the principal subspace.

Each neuron’s firing rate response to an olfactory stimulus depends on the projection to that neuron’s perceptual subspace, i.e., on the concentration of a characteristic odor blend that drives that neuron. Fixing one neuron *i*, let **u**_*i*_ = (∑ _*j*=1_ *r*_*ij*_*a*_*j*1_, …, ∑ _*j*=1_ *r*_*ij*_*a*_*jd*_) denote the *i*-th row of the matrix **RA**. The neuron’s response to an odor concentration vector **x** = (*x*_1_, …, *x*_*d*_) is given by the dot product **x** *·* **u**_**i**_. However, the projection to the perceptual subspace is given by the dot product with a unit vector **û**_**i**_ computed by normalizing **u**_*i*_. Thus, the firing rate depends on the projection to the perceptual subspace linearly, with a slope given by the *response gain*,

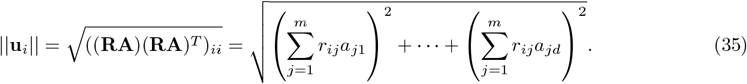

For a population of neurons, we compute the mean population response gain as 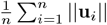.

### 3.8 Sparse odor distribution and nonlinear network

We train a feedforward network modeled on the olfactory system to decode odor concentrations sampled from the sparse odorant distribution described in Section 3.5 from receptor activation. The input is **y** = **A**(**x** + *ξ*), where **A** is the shuffled affinity matrix (data from [43]).

The network architecture consists of an input layer of 50 receptor types, 25 units corresponding to OSN types, a hidden layer with 800 units, and 110 output units. The units in both the OSN layer and the hidden layer have sigmoid activation functions with learnable gain and bias. Both layers are fully connected at initialization, weights are initialized uniformly between 0 and 1*/N*, where *N* is the number of input neurons to the layer. Weights in the receptor type to OSN type layer are constrained to be positive. Additionally, there is a *L*1 penalty applied to weights in this layer, computed as

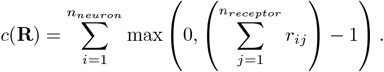

Independent neural neural noise *ζ*_*i*_ sampled from a Gaussian distribution with mean zero and variance *v*_*n*_^2^ is added inside the nonlinearity. Each neuron’s response is thus given by Sigmoid(*g*(*r*_*ij*_*y*_*j*_ + *b*_*j*_) + *ζ*_*i*_), where the sigmoid function is given by Sigmoid 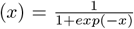 and *g*_*i*_, *b*_*i*_ are the gain and bias of neuron *i*, respectively. The output layer is linear.

We use a training set of 4 × 10^4^ samples (**y**, *v*_*e*_, *ζ*) from our olfactory signal distribution, environmental noise distribution, and neural noise distribution. We trained the network for 600 epochs using the Adam optimizer with a batch size of *B* = 256, a learning rate of 10^−3^, and exponential decay rates for first and second moments *β*_1_, *β*_2_ = .999, .999.

We perform three trials. For each trial, we generate a new training set with a different sample of odor sources, We train the network for *v*_*n*_^2^ = 0.03, 0.05, 0.1 and *v*_*e*_^2^ = 0.0, 0.5, 1.0, 1.5, 2.0. We analyze the resulting expression matrix **R** with the co-expression index used in the previous section.

### 3.9 Structure of natural stimuli

We re-analyzed data from Dweck et al, presented in Figure 1 C of [44]. We downloaded the provided data set which lists the presence or absense of 29 odorants across headspace extracts from including 34 fruits extracts, 7 microbe extracts, and 11 fecal extracts. We calculated the correlation of the presence/absence across extracts between all pairs of odorants using the python function corr.

Additionally, for each odorant, Dweck et al. list which of the six maxillary palp neurons the odorant activated in Figure 1C. While five of the neurons single-express one receptor type, the pb2A neuron co-expresses Or 33c and Or 85e. Based on analysis of chemical structure, the authors conclude that the pb2A neuron’s response to strawberry furanone is driven by Or33c, while the pB2A neuron’s response to (-)-camphor, alpha- and beta-ionone is driven by Or85e.

Based on this, we select the pairs of odorants which do not activate the same receptor. We plot the distribution of correlations of these pairs of odorants in Figure 7 B. We note that the correlation between strawberry furanone and (-) camphor, the main ligands for Or33c and Or85 in this data set, is the highest correlation between odorants which do not activate the same receptor prestent in this data set.

Note that strawberry furanone is a synonym of furaneol methylether, which is the name used in the text of [44].

## 4 Supplementary figures

**Fig. S1.**
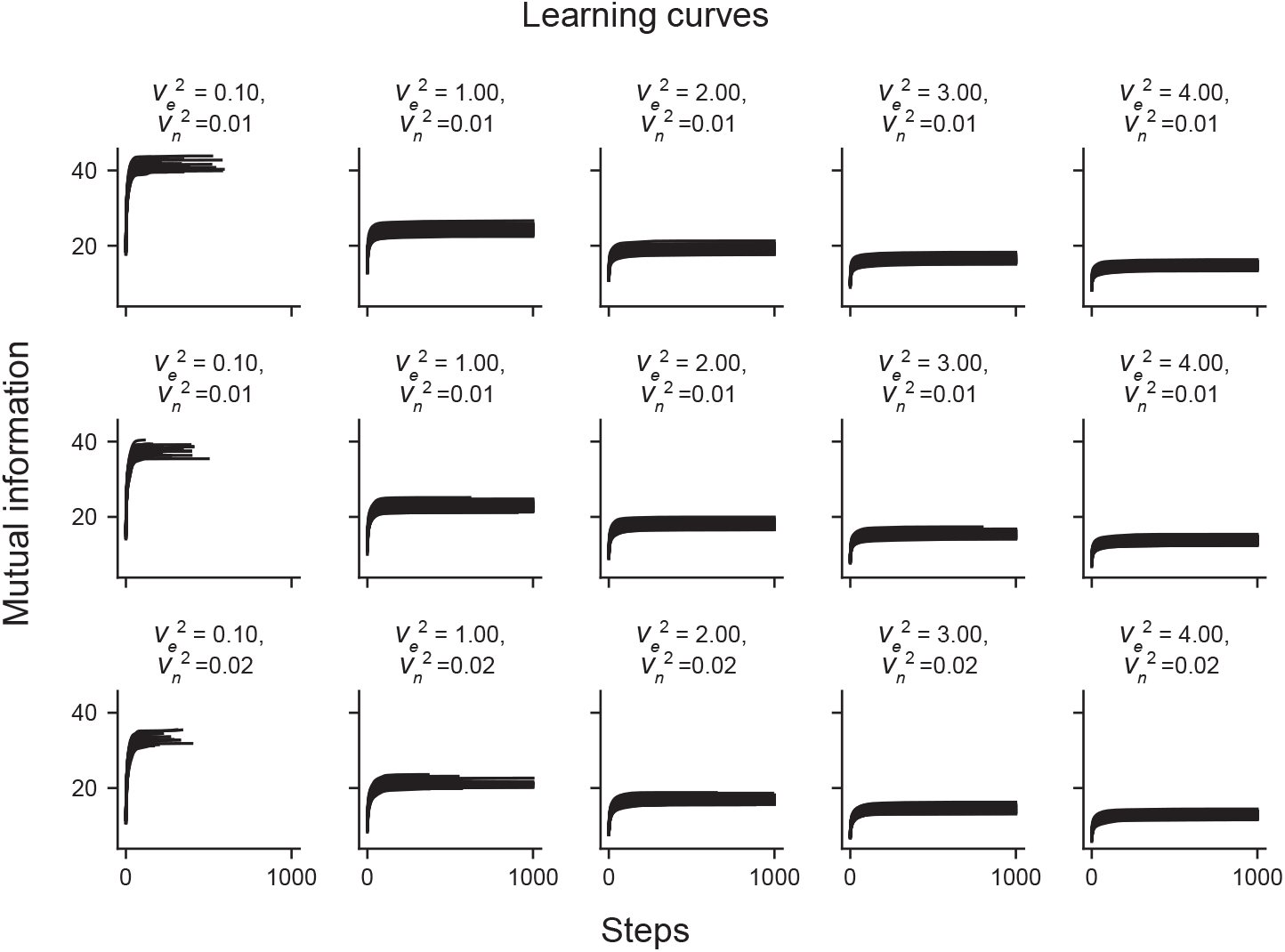
Learning curves for the optimization corresponding to Fig. 5. Mutual information vs. optimization step for fifty trials for each level of environmental noise 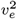 (columns) and neural noise 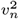 (rows). Some curves terminate early when gradient condition is met.

**Fig. S2.**
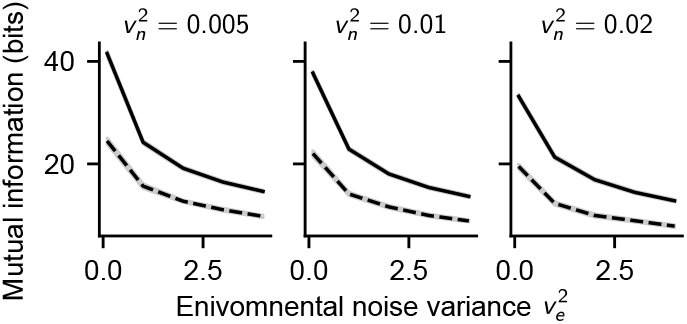
Mutual information from optimized co-expression matrix (solid line) and shuffled co-expression matrix (dashed line) as a function of environmental noise level *v*_*e*_^2^. Corresponds to Fig. 5E.

**Fig. S3.**
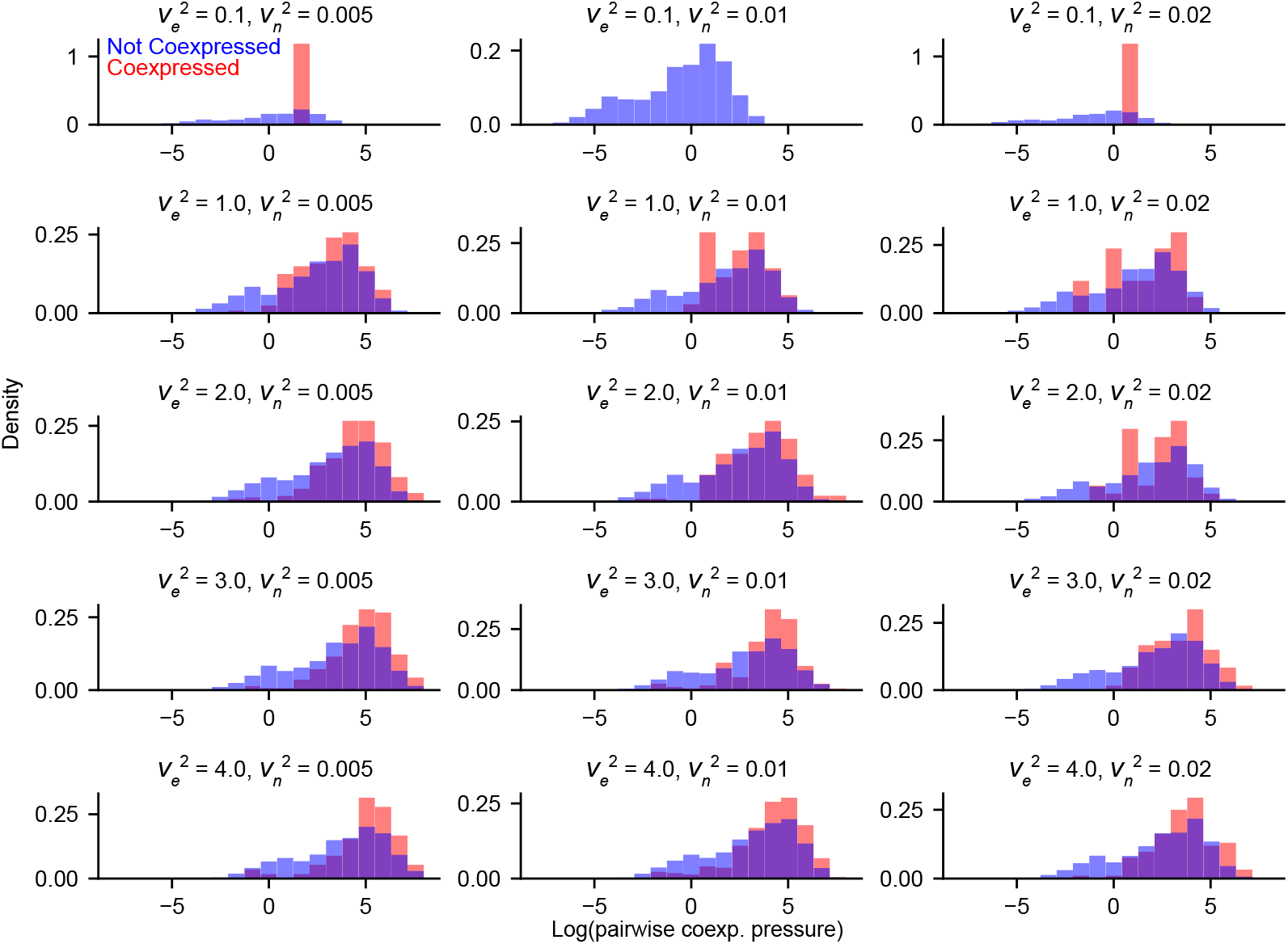
Histograms of pairwise co-expression pressure for all values of 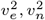. Red: co-expressed pairs of receptors. Blue: non-co-expressed pairs of receptors. Pooled over fifty trials.

**Fig. S4.**
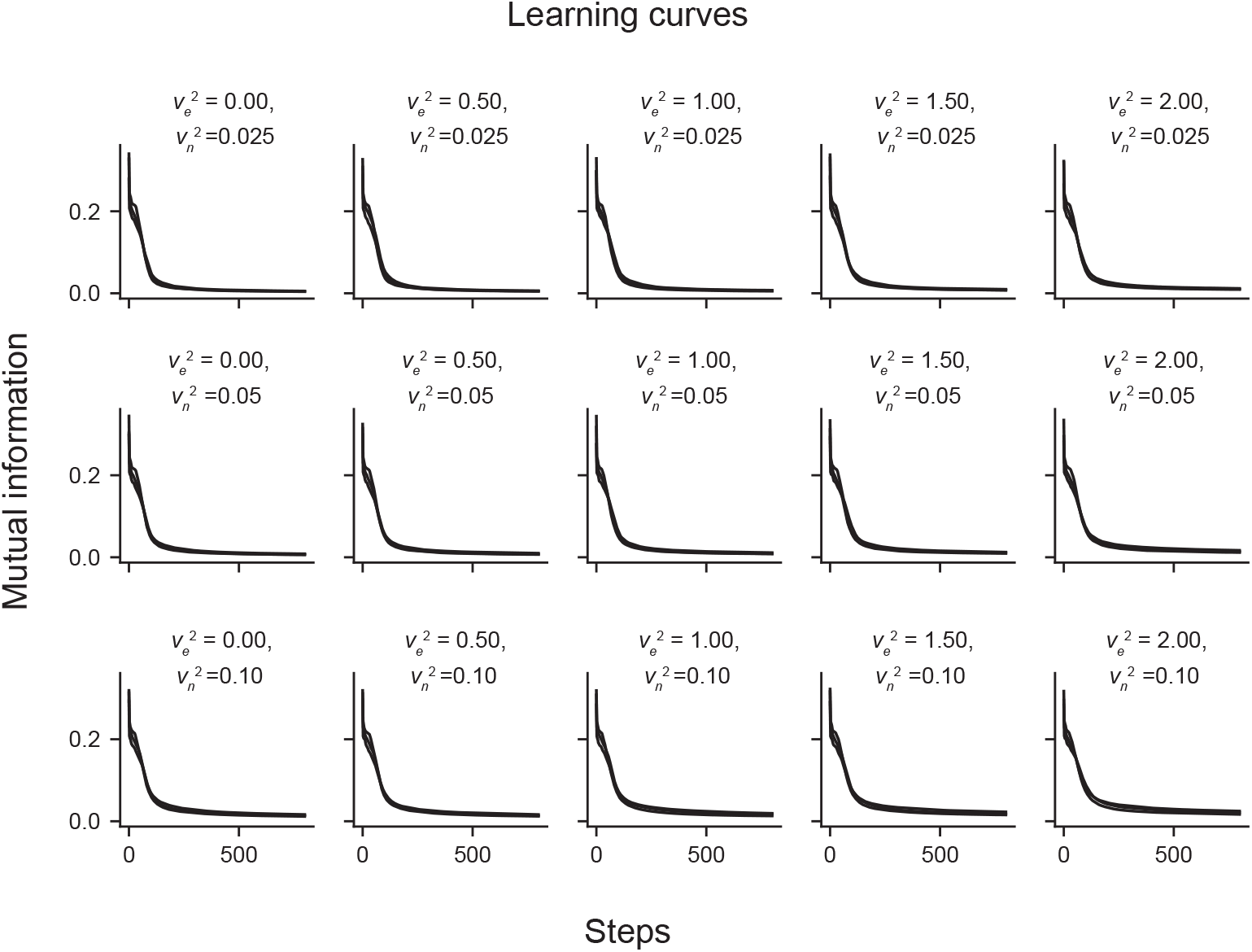
Learning curves for the optimization corresponding to Fig. 6. Mean squared error loss vs. optimization step for three trials for each level of environmental noise 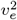 (columns) and neural noise 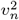 (rows).

**Fig. S5.**
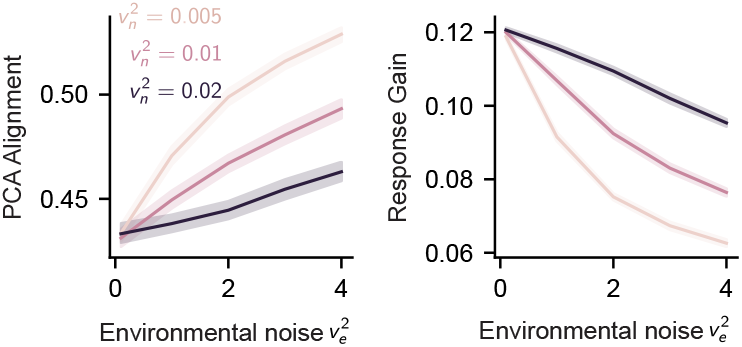
(A) PC alignment of the optimized expression matrices as a function of environmental noise variance *v*_*e*_^2^ for neural noise variance *v*_*n*_^2^ = 0.005, 0.01, 0.02 (B) Mean response gain of all neurons as a function of environmental noise variance *v*_*e*_^2^ for neural noise variance *v*_*n*_^2^ = 0.005, 0.01, 0.02. Both panels are computed from the optimized expression matrices in Fig. 5, and are averaged over fifty trials.

